# Ischemic Injury Drives Tumor Growth via Accelerated Hematopoietic Aging

**DOI:** 10.1101/2025.05.03.652034

**Authors:** Alexandra A. C. Newman, José Gabriel Barcia Durán, Richard Von Itter, Jessie M. Dalman, Brian Lim, Morgane Gourvest, Tarik Zahr, Kristin M. Wang, Tracy Zhang, Noah Albarracin, Whitney G. Rubin, Fazli K. Bozal, Michael Gildea, Coen van Solingen, Kathryn J. Moore

## Abstract

**Background:** Patients with peripheral artery disease have increased risk of cancer development. Aging-associated changes in hematopoietic stem and progenitor cells (HSPCs), including inflammation and increased myelopoiesis, are implicated in both cardiovascular disease (CVD) and cancer, but their contributions to CVD-driven tumor progression are unclear.

**Objectives:** To study cancer growth following peripheral ischemia and consequent changes within the HSPC bone marrow compartment to uncover mechanisms through which altered hematopoiesis promotes oncogenesis.

**Methods:** Mammary cancer cell (E0771) growth was monitored in C57BL/6J mice after hind limb ischemia (HLI) or sham surgery. The tumor immune microenvironment, circulatory immune cells, and HSPC compartment were assessed by flow cytometry. Next-generation single-cell RNA and ATAC sequencing of HSPCs was performed to assess transcriptomic and epigenetic changes. The functional impact on tumor progression and persistence of ischemia-induced epigenetic reprogramming of HSPCs and their myeloid progeny was examined by bone marrow transplantation.

**Results:** Peripheral ischemia increased monocyte and neutrophil output at the expense of lymphocytes, driven by a shift toward CD150^hi^ myeloid-biased hematopoietic stem cells (HSCs). This was associated with accelerated breast cancer growth and increased accumulation of tumoral immunosuppressive regulatory T cells and monocytes. Increased myelopoiesis was also supported by multiomic analyses showing HLI-induced transcriptional and epigenetic enrichment for inflammatory (NLRP3 inflammasome) and aging-associated (Neogenin-1, Thrombospondin-1) signatures in subsets of monocyte/dendritic progenitors. HLI-accelerated tumor growth and myeloid-skewing was transmissible via bone marrow transplantation, indicating long-term reprogramming of innate immune responses.

**Conclusions:** Peripheral ischemia promotes inflammaging of HSCs and long-lasting alterations to anti-tumoral immunity, accelerating breast tumor growth.

## Introduction

Cardiovascular disease (CVD) and cancer are the two leading causes of death globally (1). Evidence increasingly suggests that these diseases share common risk factors and pathophysiological mechanisms in mice and humans, resulting in bidirectional cross-disease communication (2). Epidemiological studies have established that adult cancer survivors have increased risk of CVD (3), and in turn, individuals diagnosed with CVD have an increased risk of incident cancer and metastasis (4,5). The intricacies of this detrimental bidirectional relationship are not well-defined.

Clinical findings suggest that myocardial infarction (MI), a common consequence of atherosclerotic CVD, can increase cancer risk and worsen outcomes in cancer patients. These reports are supported by preclinical animal studies showing that acute MI or heart failure accelerates tumor growth in breast, lung and colorectal cancer (6–8). Several of these studies report changes in the tumor immune microenvironment, suggesting that CVD may impact anti-tumoral immunity (7). Tumors are populated by both innate and adaptive immune cells that can take on either tumor-suppressive (e.g., natural killer cells, NK; CD8+ T cells) or tumor-promoting (e.g., myeloid derived suppressor cells, MDSC; regulatory T cells, Treg) phenotypes. Innate immune cells, in particular, can be polarized to phenotypes that suppress cytotoxic cellular responses, thereby enabling tumor growth.

Recently, we demonstrated that acute MI induced immunosuppressive reprogramming of monocytes in the bone marrow, as well as circulating Ly6C^hi^ monocytes and intra-tumoral MDSCs (7). These findings are consistent with the sequelae of emergency myelopoiesis, which has been associated with accelerated tumor growth (9). Whether these changes are driven by cardiac injury or ischemic insult, however, remains unclear. The importance of elucidating this distinction becomes increasingly apparent when considering the widespread prevalence of other common sequelae of atherosclerosis: peripheral arterial disease (PAD), for instance, leads to both acute and chronic ischemia of the extremities and has been linked to increased cancer risk (10). Despite the widespread prevalence of PAD and the high mortality rate (11) associated with this pathology, only those patients with symptomatic disease are diagnostically evaluated, resulting in probable underdiagnosis of this disease (12). Though atherosclerosis has generally been associated with increased myelopoiesis (13), PAD patients in particular have been shown to demonstrate monocytosis compared to age-matched healthy controls(14), which is especially interesting when considered in conjunction with the inverse correlation between myeloid output and cancer-specific outcomes (15).

Recent studies suggest that systemic inflammatory insults can lead to long-lasting adaptations in hematopoietic stem and progenitor cells (HSPCs) in the bone marrow associated with innate immune memory. This is based on epigenetic changes in HSPCs that are maintained in their myeloid progeny and confer altered responsiveness to subsequent immune challenges (16). This heterologous innate immune memory engenders sustained myeloid skewing, which is also a prominent feature of aging (17). Indeed, self-renewing hematopoietic stem cells (HSCs) in both humans and mice exhibit a shift from balanced output of lymphoid and myeloid cells (bal-HSC) to primarily myeloid-biased output (mye-HSC) with increased age (18–20). This process is believed to be central to the diminution of adaptive immunity and the augmentation of myelopoiesis and inflammation that leads to a predisposition to developing age-related pathologies, including cancer.

In this study, we evaluate the impact of peripheral ischemia on breast tumor growth in mice and explore the construct that reprogramming of hematopoietic progenitors in the bone marrow provides a mechanistic link to altered anti-tumoral immunity post-ischemia. We hypothesize that ischemia-driven alterations in bone marrow progenitors leads to a maladaptive anti-tumorigenic immune response that is permissive to cancer progression. Our present findings show that PAD, as modeled by femoral artery ligation-induced hind limb ischemia (HLI) in mice, accelerated mammary cancer growth and altered the tumor immune microenvironment by enriching for Ly6C^hi^ monocytic cells and immunosuppressive CD4^+^ Tregs. This was paralleled by persistent changes in the circulating leukocyte pool after HLI, with preferential mobilization of myeloid cells from the bone marrow that was associated with an increased frequency of myeloid-biased HSCs (mye-HSC) relative to those with balanced output (bal-HSC). Underlying these changes, we observed that HLI induced alterations of chromatin accessibility and transcriptional profile in myeloid-derived progenitors consistent with the acquisition of an inflammatory and aged state that is associated with cancer progression (19–21).

## Methods

Extended methods included in Data Supplement

### Mouse studies

Mouse studies were in accordance with the US Department of Agriculture Animal Welfare Act and the US Public Health Service Policy on Humane Care and Use of Laboratory Animals and approved by the Institutional Animal Care and Use Committee at the NYU Grossman School of Medicine. Female C57BL/6J mice (Jackson Laboratory, stock # 00064) aged 10-12 weeks were used in all studies.

### Tumor cell handling and implantation

Mouse E0771 mammary tumor cells were maintained in RPMI 1640 (ThermoFisher, MT10040CV), 10% fetal bovine serum (Life Technologies, 10082147), and 1% penicillin/streptomycin (ThermoFisher, 15-140-122) at 37°C with 5% CO_2_, and split at 70-80% confluence. For tumor inoculation, mice were anesthetized with 1.5% isoflurane and O_2_, and 2×10^5^ E0771 tumor cells in 100uL of sterile calcium- and magnesium-free PBS (ThermoFisher, 14190144) were orthotopically implanted into the 4^th^ right mammary fat pad. Tumors were measured using a digital caliper twice a week from day 10, when tumors first become palpable, until study endpoint. Tumor volume was calculated as follows: *V* = 0.5 ∗ (*L* ∗ *W*^2^). Images of tumors were taken with an iPhone using iOS software.

### Femoral artery ligation

Three days after tumor inoculation, mice were randomly assigned to hind limb ischemia (HLI) or sham-FAL surgery. Power analysis indicated sample size equal n of at least 6 in 2-3 independent experiments. Mice were anesthetized with ketamine (100 mg/kg) and xylazine (8 mg/kg) in sterile saline via intraperitoneal injection. The left leg was shaved and a 1cm incision slightly below and parallel to the left inguinal ligament was made followed by blunt dissection of fat and soft tissues to expose the femoral artery and vein. After careful separation of nearby nerves, an 8-0 suture was used to ligate the femoral artery and vein, which prevents the vein from becoming atretic after ligation. Skin incision was closed with a 7-0 suture. Mice received Ethiqa XR (3.25mg/kg), a long-acting analgesic following surgery. Ischemia was confirmed directly after surgery, as well as 8 and 17 days later by laser doppler imaging.

At experiment endpoint, mice were euthanized by CO_2_ and exsanguinated by cardiac puncture. Blood was processed for flow cytometry, for analysis using a Complete Blood Cell Element HT5 machine (HESKA), or for plasma collection. Organs were weighed and processed for flow cytometry, RNA analysis, or paraffin embedded for immunohistochemistry. Bone marrow was harvested and analyzed from the left (ligated or sham-ligated) and right (contralateral control) legs separately. Mice were removed from the study based on the following exclusion criteria: 1. severe tumor ulceration, 2. missed (i.e. non-mammary) fat pad tumor injection, 3. tumor volume <100mm^3^ by day 24 after tumor injection.

### Bone marrow transplantation

Bone marrow was isolated from 4 donor FAL or 5 donor sham-FAL tumor bearing CD45.2 mice 21 days after HLI or sham surgery. Bone marrow from both legs was pooled, and 1×10^6^ cells in 100uL of sterile PBS were injected retro-orbitally into naïve female age-matched CD45.1 (B6.SJL-Ptprca Pepcb/BoyJ, stock # 002014) mice that were lethally irradiated (2 x 4.5Gy, 3 hours apart) one day prior. Antibiotic water was administered for 6 weeks during recovery, and 3 weeks later, E0771 tumor cells were orthotopically implanted and tumor volume was measured as described above. Blood chimerism was determined as the ratio of CD45.2 to CD45.1 out of the total CD45^+^ cells.

### scRNAseq and snATACseq with gene expression

Bone marrow from the left (ligated or sham-ligated) leg was isolated 9 days after surgery in E0771 inoculated mice. Single-cell suspensions were filtered through a 70-µm filter and unfractionated bone marrow cells (2×10^6^) were stained for viability (eFluor 506 - eBioscience, 65-0866-14), Lineage (FITC, BioLegend, 133302), Sca1 (BV421, BioLegend, 108120), and cKit (APC-Cy7, BioLegend, 105826) expression. Cells were sorted by fluorescence-activated cell sorting (FACS) machines (FACSAria IIu or FACS Symphony S6 SORP) equipped with a 100 µm nozzle at the NYU Cytometry & Cell Sorting Laboratory. Gating to exclude debris, doublets, and dead cells as described above. Cells were sorted as cKit+ Sca-1^+/–^ and processed for single nucleus (sn)ATACseq or single cell (sc)RNAseq by the NYU Genome Technology Center. See supplement for detailed methods and analyses.

### Statistical analysis

Outliers were determined by ROUT test. Normality was determined by Shapiro-Wilk test. Student’s t-test or Mann-Whitney U-test was used to determine statistical significance between two groups if data were distributed normally or not, respectively. One-way ANOVA was performed for three groups or more groups and Two-way ANOVA was used to assess interaction between two groups with two variables. A priori selection of Sidak’s post hoc MCT was used.

All statistical analyses were performed using GraphPad Prism or R studio. Threshold for statistical significance was p≤ 0.05.

### Data Availability

Sequencing data be deposited in the Gene Expression Omnibus hosted by the National Center for Biotechnology Information and will be made public upon acceptance of the manuscript.

## Results

### Tumor growth is accelerated in a model of peripheral ischemia

To investigate the impact of peripheral ischemia on breast cancer growth, we performed femoral artery ligation or a sham surgery as control three days after orthotopic implantation of E0771 cancer cells in the mammary fat pad of C57BL6J mice (Figure 1A). To control for the effects of surgery, a sham ligation procedure in which the femoral artery was exposed but not tied off was substituted for femoral artery ligation. Laser doppler imaging was used to confirm ischemia immediately following surgery, with an average 68% reduction in the perfusion ratio of the ligated versus unligated limb in HLI compared to sham surgery (Figures 1A and 1B). Perfusion in the ligated limb was fully restored by day 17 after HLI and was comparable to sham surgery levels (Figure 1B). Monitoring of tumor volume by caliper measurements revealed a greater than 2-fold increase in tumor growth in HLI compared to sham treated mice (Figure 1C), and a doubling of tumor burden at the study endpoint 21 days after surgery (Figure 1D). This degree of HLI-accelerated tumor growth is similar to what has been observed following coronary artery ligation in a mouse model of acute MI (7), suggesting that like cardiac ischemia, peripheral ischemic injury accelerates tumor growth.

**Figure 1.**
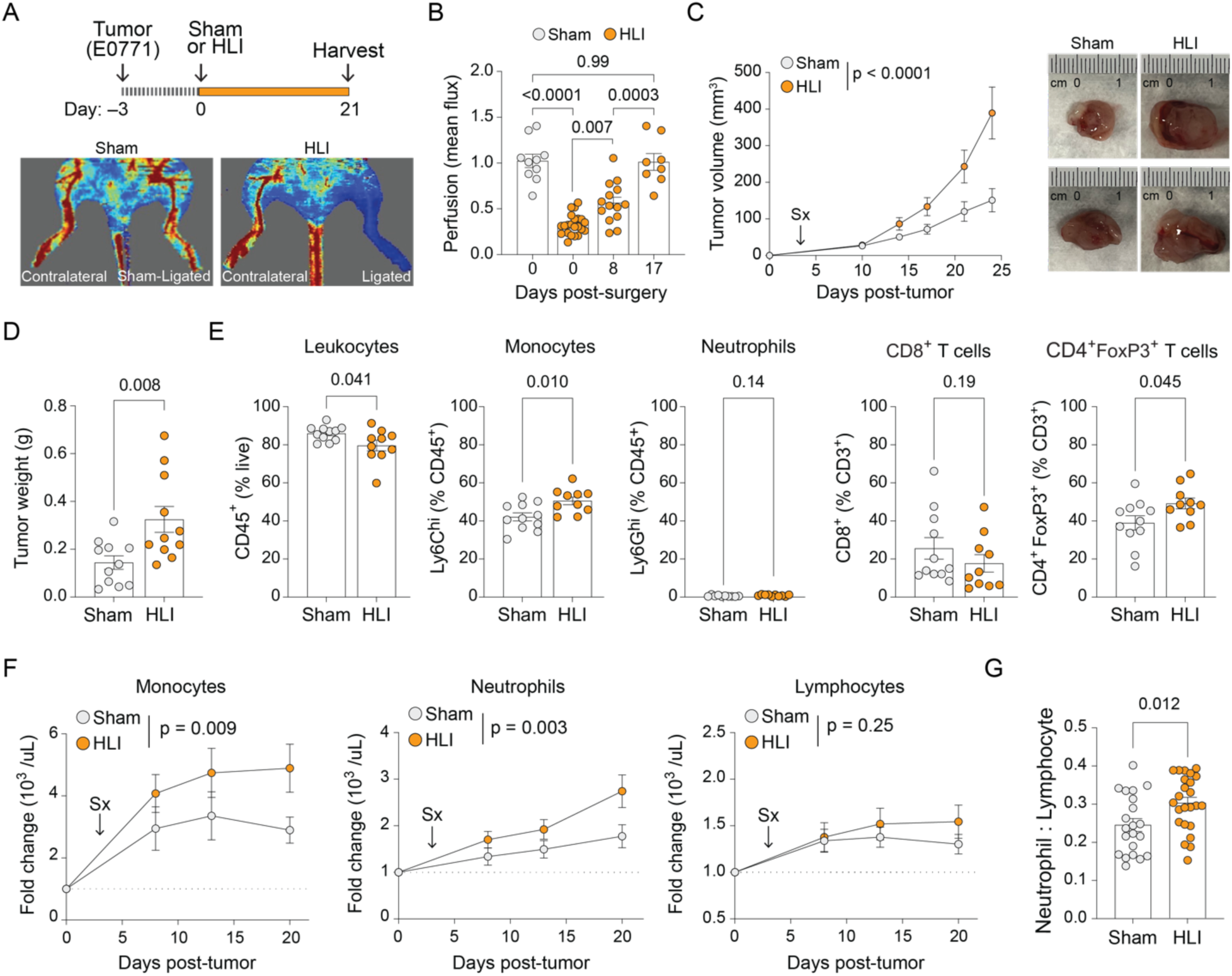
Peripheral ischemic injury accelerates tumor growth and alters immune populations. (A) Experimental design and representative laser doppler imaging of hind limb blood perfusion after femoral artery ligation or sham surgery. (B) Hind limb perfusion ratio in treated mice over time. (C) Tumor volume measured by digital caliper over the length of the study, with representative tumor images at endpoint in HLI or sham treated mice. (D) Tumor weight at endpoint. (E) Relative proportions of immune cells in tumors measured by flow cytometry. (F) Relative number of monocytes, neutrophils, and lymphocytes in the circulation of HLI or sham treated mice. (G) Neutrophil to lymphocyte ratio in the circulation at study endpoint. Data are the mean ± SEM. FC, fold change. Student’s t-test (D,E,G) or Mann-Whitney U-test (E; % Ly6G^hi^, % CD8^+^), One-way ANOVA (B) or Two-way ANOVA (C,F).

### Ischemic injury alters the tumor immune microenvironment

To evaluate the effect of peripheral ischemia on the tumor immune microenvironment, we assessed the accumulation of tumoral immune cells by flow cytometry 21 days after HLI induction (Figure 1E, Supplemental Figure 1) (7). The overall relative frequency of CD45^+^ immune cells within tumors was decreased in the HLI-experienced group compared to sham controls (Figure 1E), yet we noted an increase in the proportion of myeloid cells (CD11b^+^/CD45) within the leukocyte population (Supplemental Figure 1). While there were no changes in intratumoral Ly6G^hi^ neutrophils, we observed an increased relative frequency of Ly6C^hi^ monocytes in HLI-experienced mice (Figure 1E), which correlated positively with tumor weight (Supplemental Figure 1). Furthermore, total CD3^+^ T cells and CD8^+^ T cells tended to decrease in tumors of HLI mice, while levels of immunosuppressive CD4^+^FoxP3^+^ regulatory T cells (Tregs) were significantly increased compared to sham surgery (Figure 1E, Supplemental Figure 1). The observed enrichment in E0771 tumors of Tregs and Ly6C^hi^ monocytes, which share markers with monocytic MDSCs, is indicative of an immunosuppressive immune microenvironment that encourages tumor progression.

### Ischemia induces a systemic shift toward myeloid cells at the expense of lymphoid cells

Hind limb ischemia has previously been reported to cause a transient increase in circulating myeloid cells, which peaks at two days post-ligation, and rapidly resolves in young but not aged mice (22). Analysis of the circulating leukocyte pool in E0771 tumor bearing mice showed a sustained increase in the relative abundance of myeloid cells, including both monocytes and neutrophils, after HLI induction compared to sham surgery (Figure 1F). Additionally, the blood neutrophil to lymphocyte ratio, a negative predictor of survival in cancer (23), was elevated at study endpoint in HLI-experienced mice compared to sham surgery controls (Figure 1G). These findings indicate that peripheral ischemia in the context of tumor induces myelopoiesis and a sustained myeloid content in the blood, which is correlated with poor clinical outcomes in a variety of cancers (24,25).

### Ischemic injury mobilizes myeloid cells from the bone marrow

Myeloid cells can be mobilized from the bone marrow as well as the spleen, which is a site of extramedullary hematopoiesis, in response to infection or inflammatory stimuli. To ascertain the source of the myelopoiesis observed in HLI-experienced tumor bearing mice, we measured Ly6C^hi^ monocytes and Ly6G^hi^ neutrophils in the blood, bone marrow, and the spleen two days after surgery. Both the absolute numbers and relative frequencies of Ly6C^hi^ monocytes and Ly6G^hi^ neutrophils were increased in the blood (Figures 2A and 2B) and decreased in number in the bone marrow but not the spleen (Supplemental Figure 2) 48 hours after HLI compared to sham surgery. Mobilization of Ly6C^hi^ monocytes and Ly6G^hi^ neutrophils from the bone marrow reservoir of HLI-experienced mice was two- and three-times greater, respectively, than sham surgery controls (Figure 2C). By comparison, the relative movement of Ly6C^hi^ monocytes and Ly6G^hi^ neutrophils from the spleen was not significantly different between the HLI and sham group (Figure 2C), indicating that peripheral ischemia prompts the preferential release of myeloid cells from the bone marrow.

**Figure 2.**
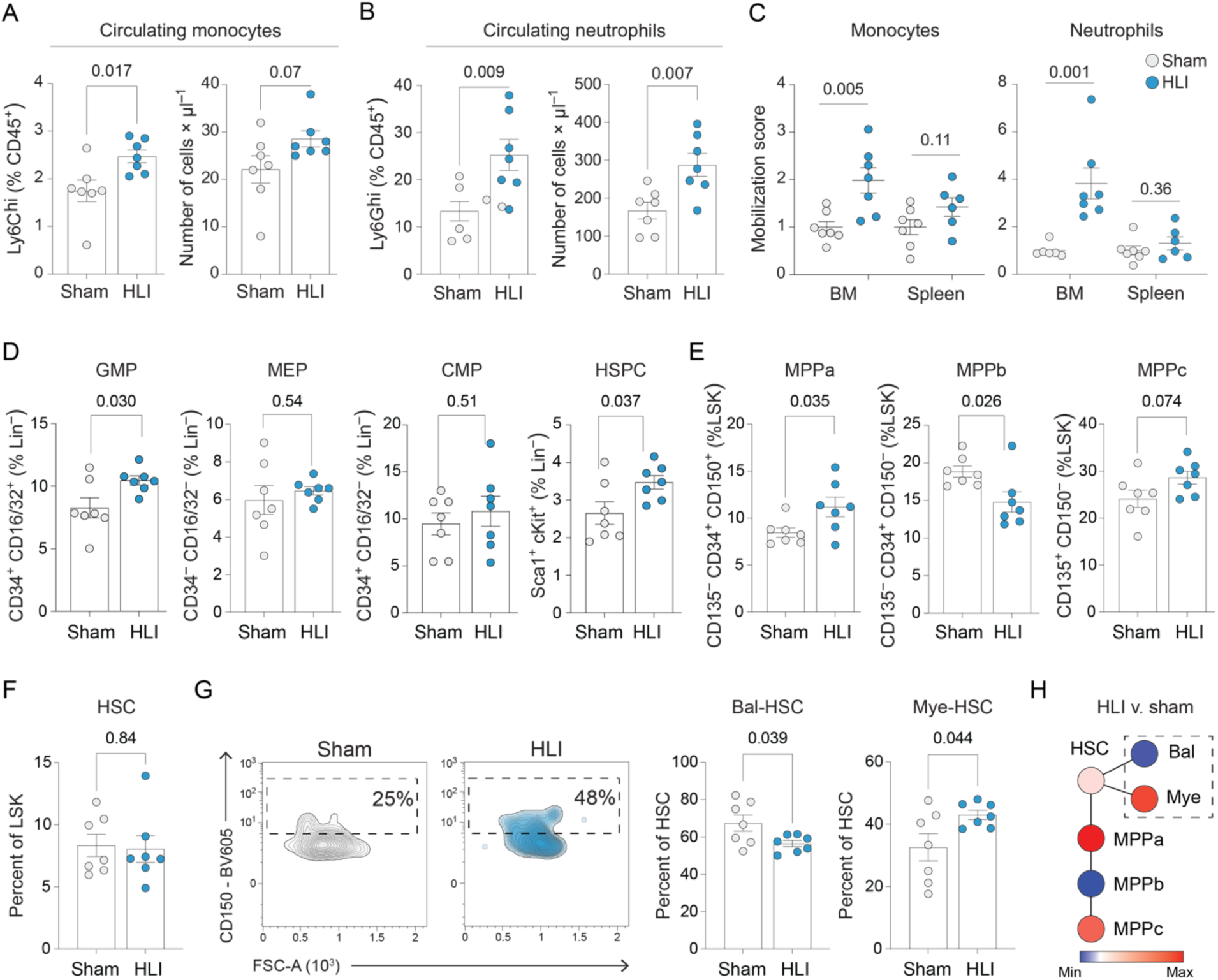
Ischemic injury increases myeloid output from the bone marrow. (A-B) Relative proportions (left) and absolute number (right) of (A) Ly6C^hi^ monocytes and (B) Ly6G^hi^ neutrophils in the blood two days after FAL or sham surgery. (C) Mobilization of monocytes and neutrophils from the bone marrow and spleen to peripheral blood. (D) Frequencies of myeloid (GMP, granulocyte-macrophage progenitor; MEP, megakaryocyte-erythroid progenitor; CMP, common myeloid progenitor) and multipotent progenitor cells (HSPC, hematopoietic stem and progenitor cell) within the bone marrow of the treated (ligated or sham-ligated) leg two days after FAL or sham surgery. (E) Frequency of multipotent progenitors (MPP) with the bone marrow. (F) Frequency of total hematopoietic stem cells (HSC) and (G) distribution of balanced and myeloid-biased HSCs gated on CD150 expression. (H) Heatmap of normalized frequencies of bone marrow hematopoietic progenitors in the ischemic leg two days after FAL or sham surgery. Data are the mean ± SEM. Student’s t-test (A-G) or Mann-Whitney U-test (A; number of Ly6C^hi^).

In concordance with these findings, the total number of bone marrow cells, including committed or multipotent hematopoietic progenitor cells, as well as stem cells, was found to be decreased in the wake of acute peripheral ischemia (Supplemental Figure 3), an effect that was partially replicated in the spleen (Supplemental Figure 4). This loss of bone marrow cellularity persisted as long as 21 days after surgery (Supplemental Figure 3), despite restoration of perfusion secondary to collateral artery formation at day 17 (Figure 1B). Thus, ischemia induces sustained rearrangement within the bone marrow niche that is ultimately associated with the promotion of tumor growth.

### Ischemia skews hematopoietic progenitors to a myeloid bias observed in aged mice

To investigate whether peripheral ischemia in the setting of cancer induces inflammatory adaptation of HSPCs, we assessed progenitor populations in the bone marrow by flow cytometry. We observed a marked expansion of Lineage^−^ cKit^+^ Sca1^+^ (LSK) HSPCs in the bone marrow of the HLI limb (Figure 2D) two days after surgery. Indeed, we saw a similar expansion of granulocyte-macrophage progenitors (GMPs, LK CD34^+^ CD16/32^+^), but not in other committed progenitor populations such as the megakaryocyte-erythroid progenitors (MEPs, LK CD34^−^ CD16/32^−^), and common myeloid progenitors (CMPs, LK CD34^+^ CD16/32^−^; Figure 2D). Together, these findings suggest that peripheral ischemia targets the most upstream progenitor populations to encourage repopulation of bone marrow reserves depleted by ischemic insult.

HSPCs can be subclassified into lineage-uncommitted and multipotent populations that can in turn generate oligopotent effector cells and mature immune populations. Analysis of bone marrow retrieved from the HLI limb two days after surgery demonstrated an increase in frequency of upstream multipotent progenitor MPPa (LSK CD135^−^ CD150^+^ CD34^+^) cells as well as MPPc (LSK CD135^+^ CD150^−^ CD34^+^) cells, which contribute primarily to lymphoid and secondarily to myeloid progenitors (Figure 2E). Notably, MPPb (LSK CD135^−^ CD150^−^ CD34^+^) cells, or the multipotent progenitor cells that predominantly give rise to myeloid cells, were decreased in the bone marrow (Figure 2E). In concert, these data suggest that peripheral ischemia induces alterations in the dynamics of myeloid and lymphoid lineage production (26).

Increased myelopoiesis associated with aging has been shown to result from a shift in the relative frequencies of HSCs with balanced and myeloid-biased output(18–20). While the frequency of HSCs (LSK CD135^−^ CD150+ CD34^−^; Figure 2F) remained unchanged between HLI and sham mice, measurements of CD150 expression, a marker used to separate bal-HSC and mye-HSCs, demonstrated that that HLI increased the proportion of mye-HSCs (CD150^hi^) and reduced the proportion of bal-HSC (CD150^lo^) in the bone marrow (Figure 2G and 2H) and the spleen (Supplemental Figure 4). These findings indicate that HLI in E0771-tumor bearing mice induces preferential differentiation of hematopoietic progenitors toward the myeloid lineage at the expense of lymphoid output (Figure 2G), a pro-inflammatory phenotype associated with aging of the hematopoietic niche (9,20).

### Ischemic injury promotes an aged, inflammatory phenotype in hematopoietic progenitors

To study the mechanisms underlying these changes to hematopoietic populations within the bone marrow in the setting of peripheral ischemia and cancer, we performed single-cell RNA sequencing (scRNA-seq) on bone marrow HSCs as well as committed and multipotent progenitors from E0771-inoculated mice nine days after HLI or sham surgery. Following unbiased Louvain clustering of 23,558 cells, we identified four lineage-uncommitted populations, as well as 11 myeloid, six lymphoid, and four megakaryocytic/erythroid progenitor (MEP) clusters (Figure 3A), in agreement with previous work (27–29). *Hlf* expression was used to identify the most immature cluster, designated as HSPCs, which is also consistent with prior literature(29). In addition to CMPs and GMPs, two committed monocyte/dendritic cell progenitor (MDP) clusters as well as eosinophil and basophil progenitor clusters were identified among the committed myeloid cell populations (Figure 3A). Consistent with our flow cytometry measurements, myeloid progenitors were expanded after HLI surgery, while lymphoid progenitors decreased in number (Figure 3B). Although the proportion of proliferating *Hlf*^+^ HSPCs was not significantly increased by HLI (Supplemental Figure 5), transcriptional proliferation signatures were significantly elevated in lymphoid-biased multipotent progenitors (*Flt3*^+^ MPPs) and *Ms4a1*^+^ CLPs in HLI-experienced mice compared to sham controls (Supplemental Figure 5). Particularly when considered alongside the observed reduction in frequency of circulating lymphocytes (Supplemental Figure 1), the induction of a transcriptional program consistent with increased proliferation among lymphoid progenitors in the bone marrow suggests that HLI disrupts normal lymphopoiesis, promoting progenitor expansion while impairing their differentiation into mature lymphoid cells.

**Figure 3.**
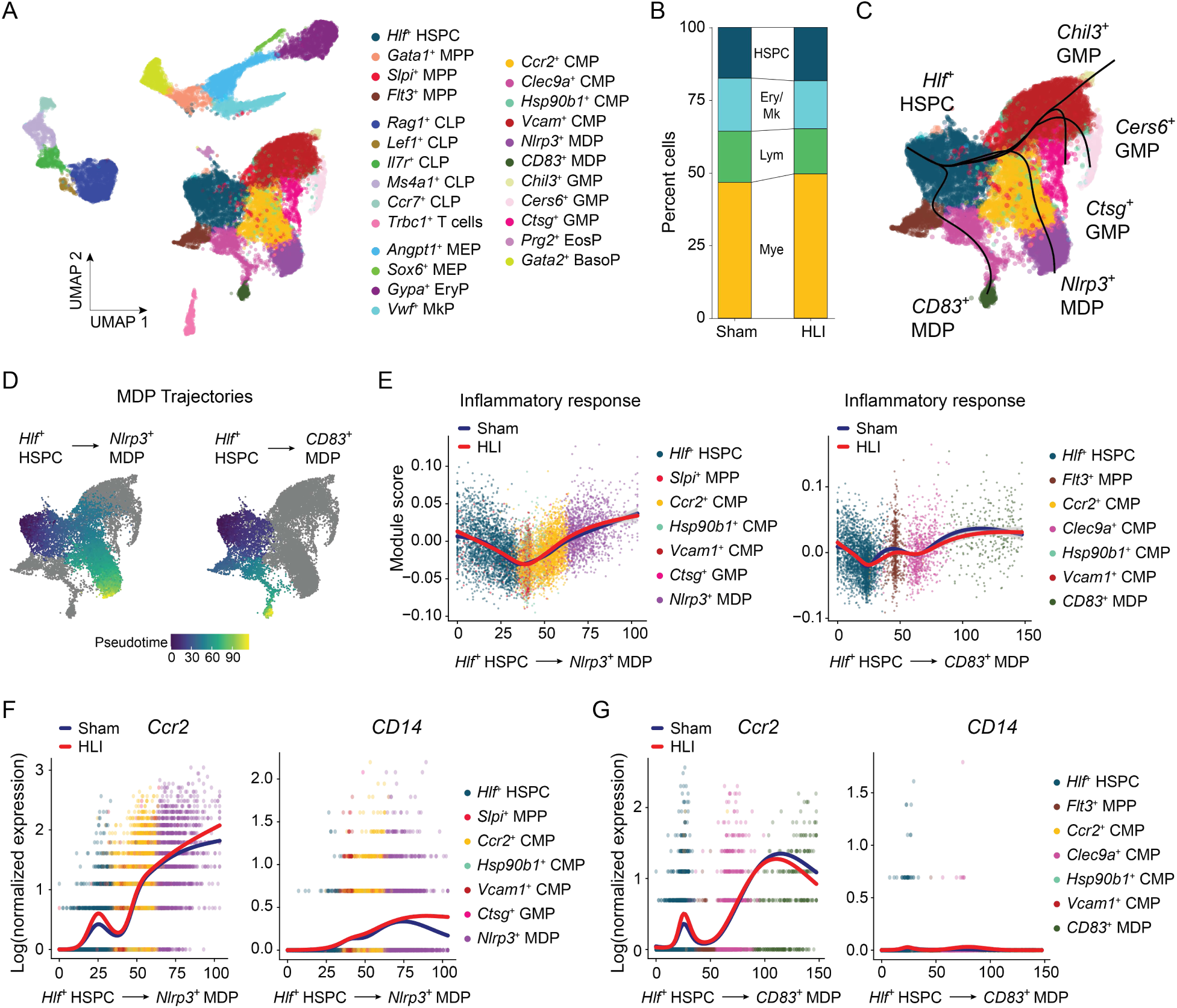
HLI induces pro-myelopoietic, inflammatory transcriptional program in the bone marrow. (A) UMAP of scRNA-seq data clustered by cell type. (B) Proportion of major cell identities, stratified by treatment. (C) UMAP showing myeloid differentiation trajectories originating from the *Hlf*^+^ HSPC cluster. (D-G) UMAP (D), hallmark gene set enrichment analysis of inflammatory pathways (E), and transcript expression (F-G) of canonical monocyte marker genes along *Nlrp3*^+^ (F) and *CD83*^+^ (G) MDP trajectories in pseudotime.

To capture the continuum of transitions and branching hierarchies of the sequenced hematopoietic cells, we reconstructed their differentiation trajectories in an unsupervised manner using cell lineage inference tool Slingshot (30). The earliest progenitors, *Hlf*^+^ HSPCs, gave rise to a single CLP trajectory (Supplemental Figure 5) and five distinct myeloid GMP or MDP fates (Figure 3C) in our model, highlighting the diversity of bone marrow myelopoietic activity. One of the two MDP clusters was distinguished by unique expression of the inflammasome component *Nlrp3* and demonstrated increasing enrichment of the “Inflammatory response” hallmark term (31) from the murine Molecular Signatures Database along its differentiation trajectory, as identified by gene set enrichment analysis (GSEA)(32,33) (Figures 3D and 3E). In addition, hematopoietic progenitors exhibited increasing expression of *Ccr2* and *Cd14* as *Hlf*^+^ HSPCs transition toward the *Nlrp3*^+^ rather than the *CD83*^+^ MDP fate, with more pronounced expression following HLI compared to sham surgery (Figure 3F and 3G). As Ccr2 is a well-established marker of pro-inflammatory monocytes originating from the bone marrow (34), these findings suggest that *Nlrp3*^+^MDPs likely give rise to inflammatory monocytes, and this is enhanced in the context of ischemic injury.

We also observed increasing expression of inflammatory gene programs along the lymphopoietic trajectory, which culminated in *Ccr7*^+^ CLP fate acquisition (Supplemental Figure 5), as well as along the myeloid path toward *Chil3*^+^ GMPs, but not *Cers6*^+^ or *Ctsg*^+^ GMPs, (Supplemental Figure 6). However, in contrast to the *Nlrp3*^+^ MDP trajectory, these programs were not differentially regulated by HLI, suggesting that in CLPs or GMPs trajectories, inflammatory gene programs are developmentally programmed rather than induced by ischemic injury.

An exacerbated inflammatory response can potentiate hematopoietic aging in the setting of cancer(9) where myeloid-biased hematopoiesis, in particular, is known to promote a feed-forward loop that further accelerates the hematopoietic aging process (17). Differential expression analyses using fitted regression model tradeSeq (35) showed that *Nlrp3* expression was increased along the differentiation path from *Hlf*^+^ HSPC to *Nlrp3*^+^ MDP fate following HLI compared to sham surgery (Figures 4A and 4B). Increased *Nlrp3* expression is associated with priming of the NLRP3-inflammasome, which drives inflammation and promotes aging by triggering the production of cytokines like interleukin (IL)-1β and IL-18 (36). Trajectory analysis showed that HLI induced a spike in *Nlrp3* transcript expression in *Hlf*^+^ HSPCs that remained elevated in *Ccr2*^+^ CMPs and *Nlrp3*^+^ MDPs (Figures 4A and 4B). In addition, in response to HLI, *Thbs1* was elevated in *Vcam1*^+^ CMPs, the population that contributed to both MDP and GMP lineages, and its expression increased again in the *Nlrp3*^+^ MDP population (Figures 4A and 4C, Supplemental Figure 6). *Thbs1* has been associated with HSC inflammaging and its deletion preserves hematopoietic health over time (37). Myosin light chain gene *Myl10*, which, along with *Thbs1*, is linked to HSPC loss (38), is also upregulated earlier along the *Nlrp3*^+^ MDP trajectory by HLI compared to sham surgery (Figure 4D). Additionally, HLI induced *Fgd4*, the transcript encoding guanine nucleotide exchange factor Frabin, which activates Cdc42 (39) and is associated with inflammatory macrophage fate acquisition in the context of ischemic injury (40), along the *Nlrp3*^+^ MDP trajectory with HLI (Figure 4E). These findings confirm that peripheral ischemia stimulates pro-inflammatory myelopoiesis, specifically driving inflammatory monocyte production. Notably, we observe that HLI affects the earliest stages of hematopoietic differentiation in the bone marrow, leading to increase expression of genes associated with inflammation and aging.

**Figure 4.**
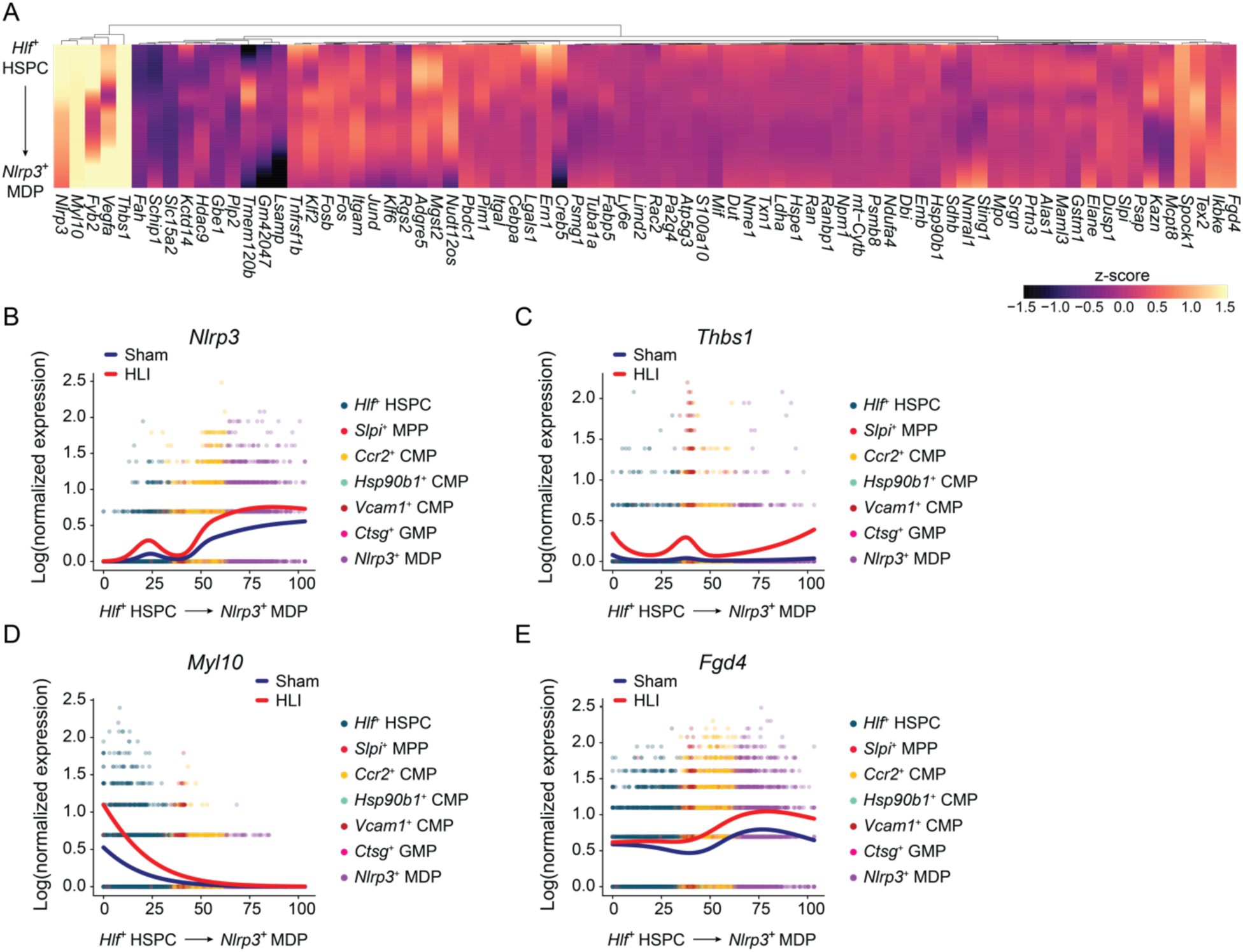
HLI induces an aged and inflammatory phenotype in myeloid progenitors in the bone marrow. (A) Heatmap depicting significant differentially expressed genes along the *Nlrp3*^+^ MDP trajectory on the y-axis. Z-score shows scaled average expression. (B-E) Transcript expression of indicated differentially expressed genes along the *Nlrp3*^+^ MDP trajectory in HLI and sham treated mice.

### Peripheral ischemia induces dysfunction in early hematopoietic bone marrow progenitors

To uncover the regulatory mechanisms driving ischemia-induced myeloid bias and their long-term effects on hematopoietic aging and inflammation, we conducted a multiomic, single-nucleus assay for transposase-accessible chromatin (ATAC)-seq paired with RNA-seq of bone marrow hematopoietic progenitors. Weighted nearest neighbor clustering identified two lineage-uncommitted clusters as well as seven myeloid, four lymphoid, and three MEP clusters (Figure 5A). Peak distribution analyses showed that compared to sham surgery, myeloid-biased progenitors after HLI carried 68% of the more accessible peaks and 57% of the less accessible peaks within the dataset (Figure 5B, left). Refinement of this analysis among myeloid progenitor populations showed that the majority of myeloid peak accessibility changes downstream of HLI occurred within an *Nlrp3*^+^ MDP cluster (Figure 5B, right). Trajectory analysis of myeloid progenitors revealed that *Nlrp3*^+^ MDPs shared an *Mgst1*^+^ precursor cluster with *CD83*^+^ MDPs (Supplemental Figure 7) and that the MDP and GMP lineages remained largely distinct, which is consistent with our findings by scRNA-seq (Figure 3). To validate these trajectories, we computed stem and proliferation “scores” as described in Giladi et al. (29), and found that as stem scores decline along the myeloid differentiation path of HSPCs, proliferation scores rise (Supplemental Figure 7), in concordance with known HSPC differentiation dynamics.

**Figure 5.**
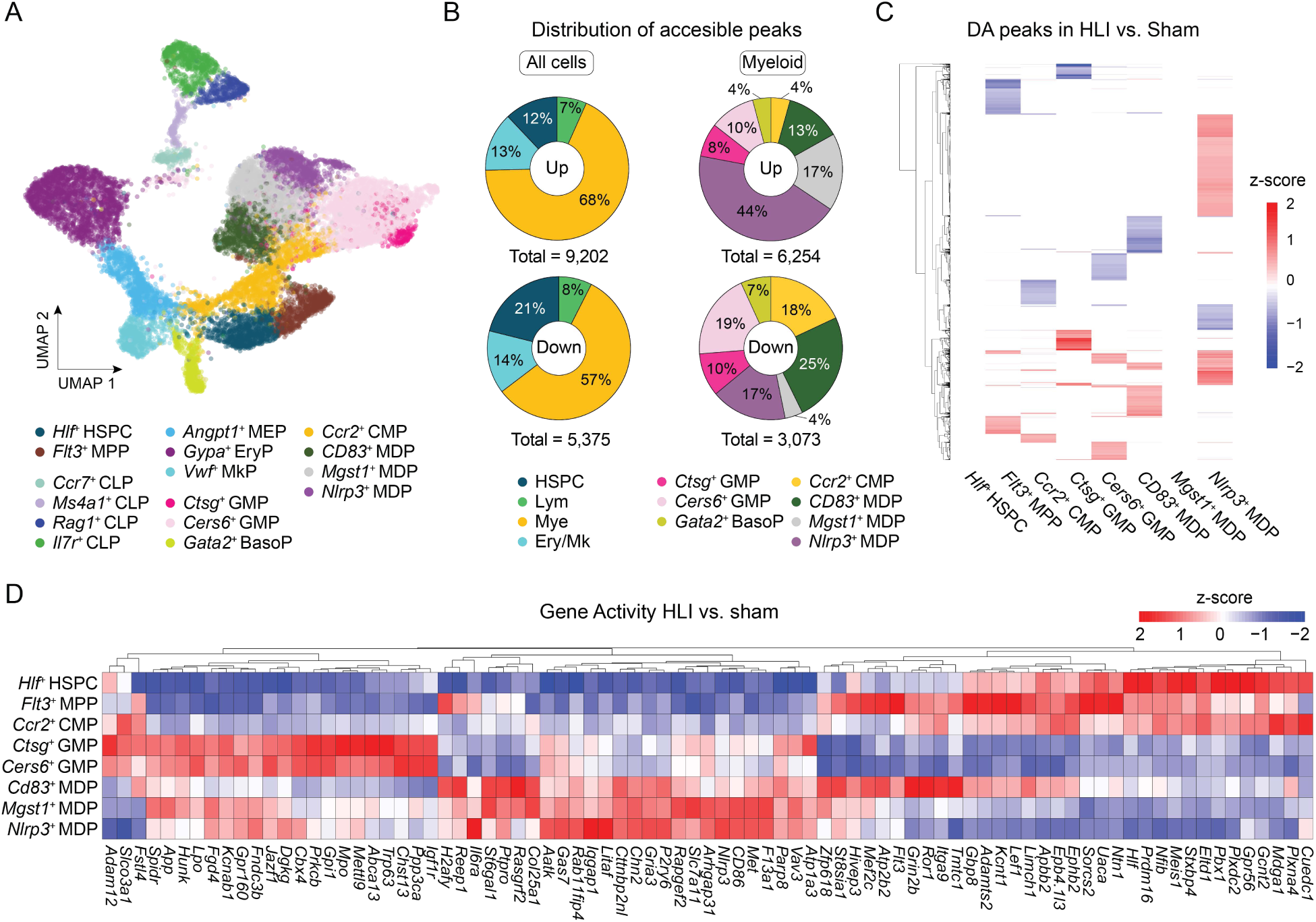
Ischemic injury promotes epigenetic reprogramming toward inflammatory state in bone marrow myeloid progenitors. (A) UMAP of single-nucleus multiomic data clustered by population. (B) Distribution of accessible peaks among major cell identities (left) and myeloid clusters (right), stratified by increased (top) or decreased (bottom) accessibility. (C) Heatmap depicting significant differentially accessible peaks in multipotent and myeloid clusters in HLI compared to sham surgery. Z-score indicates normalized read counts across peaks. (D) Heatmap depicting significant differentially active genes in multipotent and myeloid clusters in HLI compared to sham surgery. Z-score shows scaled average expression.

*Nlrp3*^+^ MDPs also displayed the majority of unique and differentially accessible peaks between HLI and sham surgery among clusters along its trajectory (Figure 5C). In general, accessible peaks encompassed more transcriptional start sites in myeloid progenitors and their more immature precursors in HLI-experienced mice (Supplemental Figure 7), suggesting enhanced transcriptional regulation. Up to 44% of these accessible regions in the myeloid clusters were located within promoter regions, which are key for gene activation (Supplemental Figure 7). Differential expression analysis of gene activity revealed increased accessibility of pro-inflammatory genes in *Nlrp3*^+^ MDPs after HLI, including *Nlrp3*, *Il6ra*, and *CD86* in *Nlrp3*^+^ MDPs (Figure 5D). These findings indicate that *Nlrp3*^+^ MDPs are a key node for ischemia-induced myeloid bias, with extensive chromatin remodeling and increased accessibility of pro-inflammatory gene loci. The changes in chromatin accessibility during differentiation show that HLI reprograms hematopoietic progenitors in the bone marrow and entrains a transcriptional and epigenetic inflammatory program linked to enhanced tumor development in their myeloid-committed progeny.

To map the specific regulatory networks driving the epigenetic changes induced by ischemia, we explored chromatin variability across the MDP clusters using transcription factor accessibility inference tool chromVAR (41). HLI resulted in increased activity at loci associated with IRF, STAT, and SPI family transcription factors, as well as PRDM1, which encodes Blimp-1, a key regulator of macrophage and dendritic cell fate (42), across all MDP clusters (Figure 6A). However, only *Nlrp3*^+^ MDPs and their precursors, *Mgst1*^+^ MDPs, showed downregulation of CEBP family loci following HLI compared to sham surgery (Figure 6A). While CEBP-α is a master regulator of steady-state granulopoiesis, CEBP-β is required for emergency granulopoiesis, facilitating rapid granulocyte mobilization amid acute inflammatory conditions (43). Conversely, IRF8 cooperates with PU.1 (encoded by *Spi1*) in phagocyte progenitors to drive monocyte production (44), while STAT2 is required for dendritic cell activation and antigen presentation (45). The concurrent reduction in *CEBP* accessibility alongside elevated *IRF*, *STAT*, and *SPI* activity—factors that promote monocyte and dendritic cell differentiation— suggests a coordinated shift away from emergency granulopoiesis toward a gene regulatory landscape favoring chronic myeloid output (Figure 6A). These transcriptional and epigenetic changes observed nine days post-HLI, both explain the myeloid expansion that follows ischemic stress and herald the more sustained, low-grade inflammation characteristic of inflammaging.

**Figure 6.**
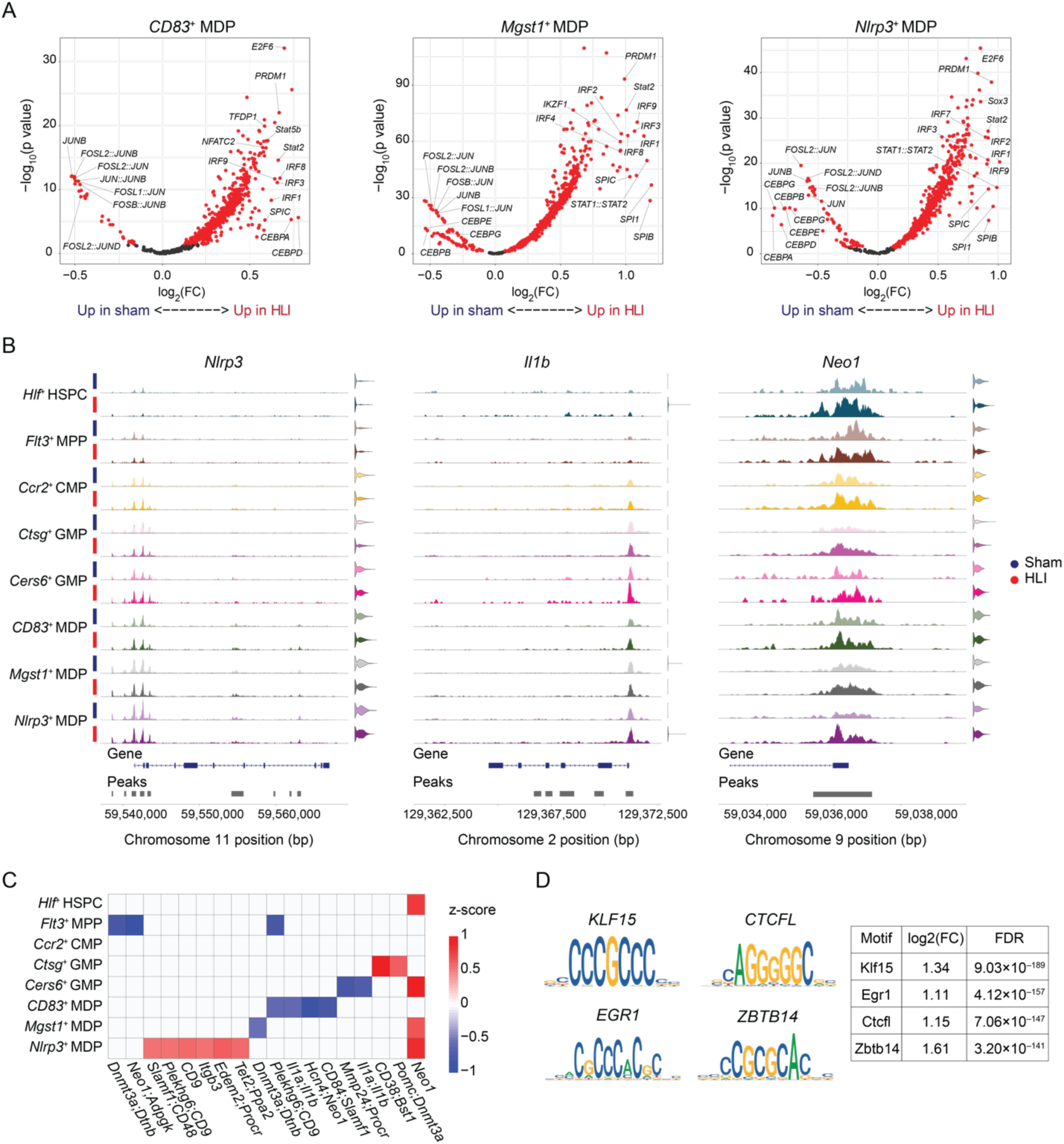
Peripheral ischemia induces inflammaging and dysfunction in myeloid bone marrow progenitors. (A) Volcano plots showing transcription factor activity in MDP clusters. FC: fold change. (B) Tracks displaying sequencing read coverage across indicated loci, stratified by treatment. (C) Heatmap depicting accessibility signal across inflammaging loci in multipotent and myeloid clusters. Z-score indicates normalized read counts across peaks. (D) Logos depicting top active transcription binding motifs in HLI compared to sham surgery (left) and quantification (right). Sequence conservation is shown with letter heights proportional to nucleotide enrichment at each position. FC: fold change. FDR: false discovery rate.

Enhanced IL-1 signaling and PU.1-driven gene programs are implicated in precocious myelopoiesis and changes associated with hematopoietic aging that promote tumor growth (46). Analysis of permissible chromatin showed that the *Il1b* locus was more accessible in *Ccr2*^+^ CMPs, *Cers6*^+^ GMPs, *CD83*^+^ MDPs, and *Nlrp3*^+^ MDPs of after HLI (Figure 6B). In addition, *Neo1*, which encodes Neogenin-1 and marks myeloid-biased HSCs in opposition to balanced HSCs (47), showed increased chromatin accessibility with HLI compared to sham surgery (Figure 6B), a phenotype associated with aging (20,48). We also observed more permissive chromatin at other inflammaging marker loci, including *CD9*, *CD48*, and *Thbs1*, particularly in *Nlrp3*^+^ MDPs (Figure 6C, Supplemental Figure 8). Of these, the *Neo1* and *Thbs1* loci were primed in *Hlf*^+^ HSPCs and maintained increased accessibility in all downstream myeloid clusters, emphasizing that the epigenetic changes brought about by ischemic injury in early hematopoietic progenitors reverberate throughout the myeloid differentiation path (Figures 6B and 6C). Motif analyses revealed that Klf15, Egr1, Ctcfl, and Zbtb14 binding sites were enriched in regions of open chromatin following HLI (Figure 6D). EGR1 regulates HSC homeostasis (49,50) and is induced downstream of Neogenin-1 signaling in aging HSCs(48), while Klf15 has been identified as a regulator of aging-related genes in the brain (51). Together, these findings suggest that ischemic injury leads to distinct epigenetic changes characteristic of inflammaging in HSCs and myeloid progenitors at various stages of differentiation that may drive inheritable, long-term immune cell reprogramming to alter cancer responses.

### Ischemia imprints persistent and transferable pro-tumorigenic changes within the bone marrow

To evaluate the longevity of the pro-tumorigenic influence of ischemia in the bone marrow, we isolated bone marrow from E0771-tumor bearing mice 21 days after HLI or sham surgery. We then transplanted the HLI-experienced and sham donor marrow (CD45.2) into naïve lethally irradiated recipient (CD45.1) mice. Following recovery and reconstitution of the bone marrow, we injected E0771 tumor cells into the mammary fat pads and followed tumor growth in the recipient mice (Figure 7A). Notably, we observed a doubling of tumor volume and tumor weight at endpoint in HLI-experienced bone marrow recipients compared to those that received sham bone marrow (Figures 7B and 7C). To exclude the possibility that altered tumor growth was due to differences in bone marrow reconstitution, we assessed CD45.1 and CD45.2 levels in the blood (Figure 7D). This analysis of peripheral chimerism showed equivalent reconstitution in transplanted mice. Furthermore, in contrast to the decrease in bone marrow cellularity noted after direct exposure to HLI surgery, no differences in cellularity were observed in bone marrow recipient mice in the HLI and sham groups (Supplemental Figure 9).

**Figure 7.**
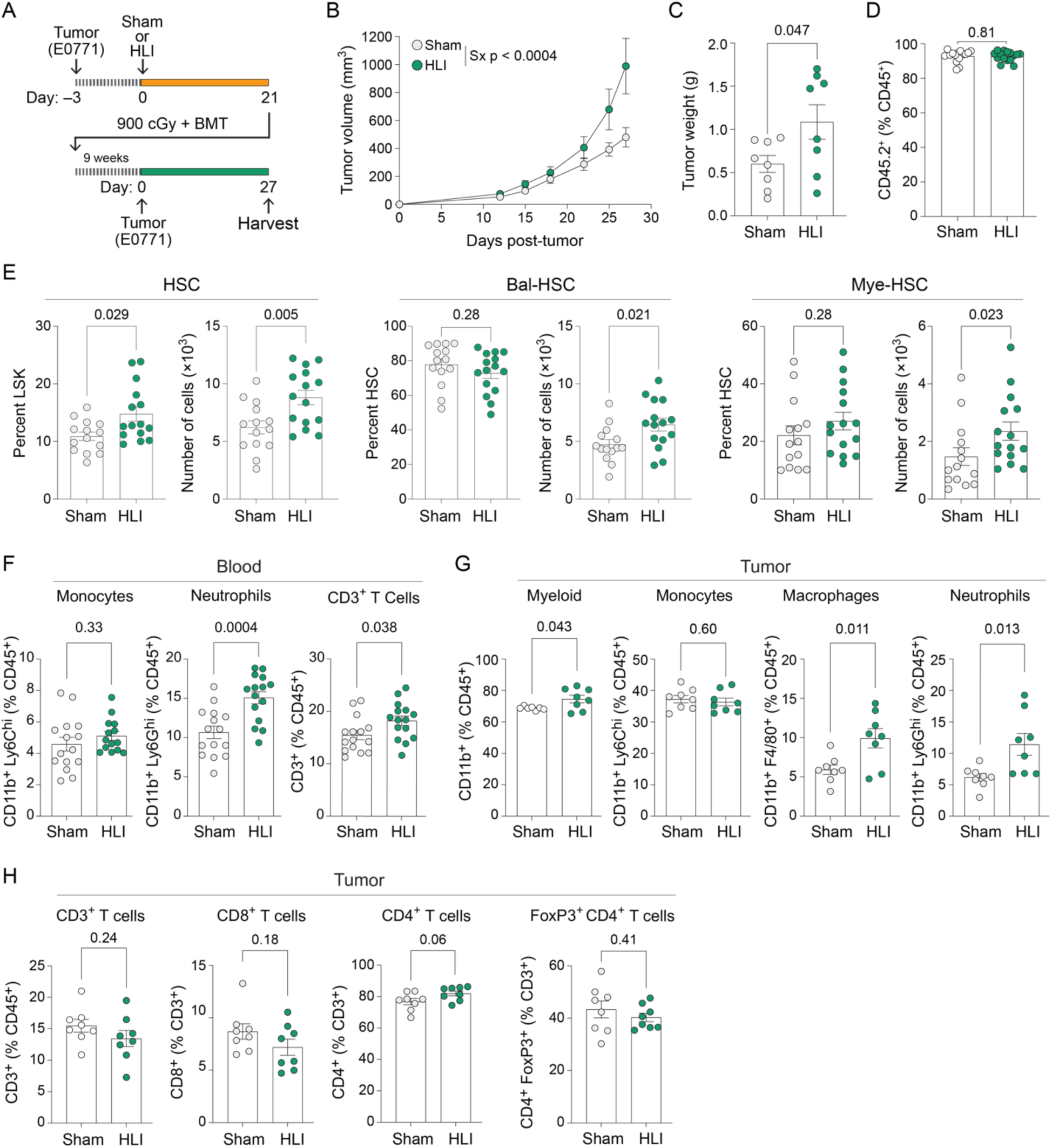
Ischemia imprints persistent and transferrable pro-tumorigenic changes within the bone marrow. (A) Experimental design of bone marrow transplantation experiments. Bone marrow was harvested from CD45.2 tumor bearing mice that received HLI or sham surgery 21 days earlier and transplanted into lethally irradiated CD45.1 recipient mice. After nine weeks recovery, E0771 mammary cancer cells were implanted into recipient mice. (B) Tumor volume over time and (C) tumor weight at endpoint in HLI and sham bone marrow recipients. (D) CD45.1/2 chimerism in the blood of CD45.1 recipient mice at time of harvest. (E) Relative proportions (left) and absolute number (right) of HSCs, balanced (Bal)-HSCs, and myeloid-biased (Mye)-HSCs in the bone marrow of recipient mice. (F) Flow cytometric analysis of Ly6C^hi^ monocytes, Ly6G^hi^ neutrophils, and CD3^+^ T cells in the blood. (G-H) Flow cytometric analysis of immune cells within tumors of recipient mice to identify (G) CD11b^+^ myeloid cells, CD11b^+^ Ly6C^hi^ monocytes, F4/80^+^ macrophages, CD11b^+^ Ly6G^hi^ neutrophils, and (H) CD4^+^ and CD8^+^ T cells, and Tregs (FoxP3^+^CD4^+^). Data are the mean ± SEM. Student’s t-test (C,D,E,F,G) or Mann-Whitney U-test (% HSC) or Two-way ANOVA (B).

Flow cytometric analysis of the bone marrow revealed an increase in total number and frequency of HSCs in HLI compared to sham bone marrow recipients (Figure 7E), despite equivalent HSC numbers in the donor marrow (Supplemental figure 3). We also observed augmented expansion of HSC content in the spleens (Supplemental Figure 10) of HLI compared to sham bone marrow recipients. However, there was no difference in the frequency of bal-HSCs and mye-HSC in either the bone marrow or the spleen (Figure 7E and Supplemental Figure 10), nor were there changes in frequency or number of other progenitor population in the bone marrow (Supplemental Figure 9). Taken together, these data suggest a selective expansion of the HSC compartment in response to chronic regenerative demand, consistent with features of hematopoietic stress and early hallmarks of HSC aging.

Analysis of mature immune cell populations in the blood revealed increases in Ly6G^hi^ neutrophils and CD3^+^ T cells, but not Ly6C^hi^ monocytes in HLI compared to sham bone marrow recipients (Figures 7F). In searching for the source of these increases, we noted elevated quantities of CD45^+^ leukocytes, including Ly6G^hi^ neutrophils and CD3^+^ T cells, as well as HSPCs, in the spleens of HLI compared to sham bone marrow recipient mice (Supplemental Figures 11). Notably, the tumor immune microenvironment of HLI-bone marrow recipient mice continued to demonstrate a myeloid bias, with significant increases in CD11b^+^ myeloid cells compared to sham-bone marrow recipients (Figure 7G). While we observed no differences in T cell frequencies (Figure 7H), there was a proportional doubling of F4/80^+^ macrophages and Ly6G^hi^ neutrophils in tumors of HLI-bone recipients (Figures 7G), suggestive of maladaptive myelopoiesis that exacerbates tumor growth.

## Discussion

In these studies, we establish that peripheral ischemia accelerates tumor growth through epigenetic reprogramming of bone marrow progenitors to an inflammatory and aged phenotype that induces long-term changes in anti-tumoral immune responses. Alongside larger tumors, mice that experienced HLI showed alterations in the tumor immune microenvironment three weeks post-ischemic injury, particularly higher numbers of Ly6C^hi^ monocytic cells and Tregs. This immunosuppressive TME was paralleled by persistent changes in the circulating leukocyte pool after HLI, with preferential mobilization of myeloid cells from the bone marrow. Analysis of bone marrow HSPCs revealed that HLI led to increased proportions of my-HSCs, a phenotype associated with aging, that results in systemic skewing of leukocytes to favor myeloid cells at the expense of lymphoid cells. This was accompanied by alterations in chromatin accessibility and gene expression within both HSPCs and lineage-committed myeloid progenitors consistent with the acquisition of an inflammatory and aged state, which has been associated with cancer progression (19–21). Notably, transplantation of HLI-trained bone marrow to naïve recipients conferred accelerated breast cancer growth with increased tumoral myeloid cell accumulation. Collectively, these results suggest that peripheral ischemia increases susceptibility to cancer growth through long-lasting inflammatory memory imprinted in myeloid cell progenitors.

### Peripheral ischemia accelerates tumor growth

The current body of evidence describing the adverse effect of cardiovascular disease or injury on oncogenesis postulates that this process is mediated several mechanisms including immune modulation, altered circulating factors and cytokines, and regulation of microbiota (6–8,52–59). Building upon prior findings by our group that cardiac ischemia induces sustained monocytosis in the setting of breast cancer and it biases Ly6C^hi^ monocytes in the bone marrow towards an immunosuppressive phenotype that accelerates tumor growth (7), our current study shows this effect to be agnostic to the location of ischemic insult. Like acute MI, HLI induces a doubling of the tumor growth rate, persistent myelopoiesis and a myeloid bias in mammary tumors alongside enrichment of immunosuppressive Tregs. These results collectively suggest that the oncogenic effect of CVD is at least partially independent of direct cardiac injury and reinforces the notion that immune modulation is a key regulator of reverse cardio-oncologic pathologies.

A coordinated innate immune response following ischemic injury is critical for tissue repair. While emergency myelopoiesis is a common response to both peripheral and cardiac ischemia, this effect is typically self-limiting, with blood monocyte levels peaking three days following the inciting event and gradually returning to homeostatic levels within a week (22,60). In the setting of nascent breast tumors, however, ischemic injury results in sustained monocytosis that persists for three weeks, suggesting an interaction between the ischemic tissue, the bone marrow and the tumor. In the case of HLI, myeloid bias in the circulation persisted even after collateral artery formation and consequent restoration of blood flow suggesting that ischemia induces long-lasting, systemic alterations to monocyte production. These cells are key regulators of the tumor microenvironment, where monocytes their differentiated progeny can assume a multitude of tumor-promoting accessory functions, including fostering tumor immune evasion and angiogenesis, as well tumor cell proliferation, migration, invasion and metastasis (24). Notably, elevated levels of circulating monocytes correlate with poor clinical outcomes in a variety of cancers (24,25).

### Ischemia accelerates aging and inflammation of bone marrow progenitors

Using flow cytometry, scRNA-seq, and snATAC/RNA-seq, we performed a detailed examination of the response of the bone marrow hematopoietic progenitor compartment after peripheral ischemia in tumor bearing mice. Our analyses reveal that ischemic injury biases the most immature cell population toward myelopoiesis while inducing inflammatory and aged epigenetic and transcriptomic changes that are associated with a cancer permissive environment. First, we noted that HLI increased the proportion of my-HSC and reduced the proportion of bal-HSC in the bone marrow and spleen. This shift from bal-HSC to my-HSCs is associated with aged immunity and increases myelopoiesis at the expense of lymphopoiesis (9,18,19). Such age-related changes in self-renewing HSCs are associated with increases in myeloid-associated inflammation and reductions in adaptive immunity, both of which are detrimental in the setting of cancer (17). Indeed, although common lymphoid progenitor (CLP) populations exhibited increased proliferation signatures after HLI, this response appears insufficient to counterbalance myeloid expansion, as their overall proportion remained lower than that of the myeloid progenitors. Second, we found that HLI-induced the most extensive changes in permissible chromatin in myeloid progenitors, with the majority occurring within an *Nlrp3*^+^ MDP cluster that showed increased inflammatory gene activity, including *Nlrp3*, *Il6ra*, and *CD86*. Indeed, complementary scRNA-seq analyses showed elevated expression of *Nlrp3* along the differentiation trajectory from *Hlf*^+^ HSPCs to *Nlrp3*^+^ MDPs, which likely give rise to Ly6C^hi^ monocytes, suggesting increased priming of the inflammasome that regulates IL-1β secretion in HLI-experienced bone marrow. Moreover, multiple myeloid progenitor populations (CMP, GMP, MDP) exhibited more permissive chromatin at the *Il1b* locus, further implicating the NLRP3-IL1 axis in the dysfunctional hematopoietic response in tumor bearing mice following HLI.

Studies have shown that myelopoiesis accelerates tumor growth through activation of IL1-IL1R axis, which simultaneously propagates hematopoietic aging (9). In HLI-experienced mice, enhanced myelopoiesis occurred alongside enrichment of EGR1 transcription factor motif accessibility in HSPCs and myeloid clusters, which is associated with HSPC aging (48,61). Notably, across multiple populations, we observed increased chromatin accessibility at aging-associated loci including Neogenin-1 (*Neo1*), a downstream target of EGR1 that has been linked to myeloid bias in HSPCs (20,48). Additionally, we detected elevated expression of Thrombospondin-1 (*Thbs1*), which has been implicated in hematopoietic aging (37), specifically within monocyte progenitor lineages. Trajectory analyses provided further insight into lineage differentiation dynamics, revealing that these inflammaging markers progressively intensified along the monocyte-producing *Nlrp3^+^ MDP* cluster. Indeed, the upregulation of both *Nlrp3* and *Thbs1* across the differentiation trajectory of *Hlf^+^* HSPCs toward *Nlrp3^+^*MDP cells implicates peripheral ischemia in hematopoietic aging.

These findings reveal that peripheral ischemic injury drives profound reprogramming of HSPC behavior—marked by inflammatory gene activation, skewed myeloid differentiation, and coordinated epigenetic remodeling—that may accelerate tumor progression. As a result, targeting inflammation may offer a promising therapeutic strategy at the intersection of cancer and cardiovascular disease. Nonetheless, further investigation is needed to clarify how ischemia reshapes the bone marrow microenvironment, including its effects on the hematopoietic stem cell niche and stromal-immune crosstalk. A deeper understanding of these mechanisms will be essential for developing interventions that restore hematopoietic balance in patients with coexisting malignancy and cardiovascular pathology.

### Study Limitations

Our findings link experimental ischemia to dysfunctional anti-tumoral immune responses and add to the growing literature supporting a reverse cardiology-oncology effect in which CVD can influence tumor growth and progression. Nevertheless, our findings have several limitations. First, our studies employ a mouse model of PAD to investigate the mechanisms by which peripheral ischemia alters growth of the mammary luminal B tumor type, E0771, as we and others have previously shown that CVD-mediated changes in immune cells dispose E0771 tumors to progress (6,7). Assessing the effect of peripheral ischemia on other cancer types is an important avenue of future study, and further studies in human samples will be required to confirm our findings in breast and other cancers. We describe epigenetic and inflammatory effects of ischemic injury on immune cell precursors within the bone marrow that predispose to hematopoietic aging in the setting of cancer. Investigations in humans will require studies of hematopoietic stem cells and their myeloid progeny following ischemic insult to establish if similar signatures of hematopoietic aging and IL-1 signaling are present that predispose to myeloid bias, and to determine how long they persist. Second, our findings reveal augmented activity of NLRP3-IL-1 along the differentiation trajectory from HSC to MDP after ischemic insult, which along with known roles for IL-1 in emergency myelopoiesis (46) and hematopoietic aging (62), implicate IL-1 as a candidate factor in HLI-induced hematopoietic dysfunction. Recent studies in aged mice showed that disrupting IL-1 receptor signaling could normalize hematopoietic aging and slow the growth of solid tumors (9); similar experiments could be performed in the context of HLI to understand the contribution of IL-1 signaling to ischemia-induced myeloid bias and accelerated tumor growth. However, it is also possible that IL-1 may also cooperate with other inflammatory factors, and this remains to be investigated. Third, the scope of our studies is limited to elucidating the effects of ischemia on immune cells and their precursors rather than circulating factors, although both have been shown to affect cancer progression (6–8,52–59). As it is most probable that a combination of factors contributes to tumor growth downstream of CVD, understanding this multimodal interplay is highly relevant. Further study is warranted to evaluate the potential influence of circulating factors related to muscle repair processes after peripheral ischemia on tumor growth. Finally, despite the transference of the HLI-induced protumorigenic immune phenotype by bone marrow transplantation, the underlying epigenetic changes that confer this maladaptive response and how long they persist for, remain to be addressed.

### Conclusions

These studies demonstrate that peripheral ischemia results in stable imprinting of early hematopoietic progenitors to skew immune output to a myeloid bias, accelerating mammary tumor growth.

### Clinical Perspectives

Peripheral arterial disease is one of the most frequent sequelae of atherosclerotic cardiovascular disease, affecting approximately two hundred million individuals worldwide. Patients typically present with intermittent claudication, or pain during ambulation that resolves with rest. While these patients are routinely assessed for other cardiovascular diseases, such as coronary artery disease, current treatment paradigms do not consider the increased risk of cancer development within this population. Indeed, the data presented here indicate that patients experiencing intermittent claudication could benefit from additional cancer-specific testing to account for the heightened vulnerability to oncogenesis conferred by this comorbidity. Furthermore, clinicians do not generally screen for PAD in those without symptoms, who constitutes an estimated 50% of the patient population (63). The results discussed in this study, however, suggest potential merits to the implementation of routine screening protocols to evaluate for the presence of PAD and subsequently augmented cancer screening guidelines for patients who are diagnosed. Consider, for instance, current breast screening protocols: women at average risk are encouraged to receive a mammogram every other year from the ages of 40 through 74. The increased breast cancer risk associated with diagnosis of PAD, and the high prevalence of this diagnosis in older adults, however, could suggest some benefit to extending the upper limit of this threshold. While such concepts would need to be evaluated in rigorous clinical trials before being utilized in clinic, these efforts could stand to benefit hundreds of millions of patients affected by peripheral ischemia who may consequently carry increased risk of developing certain cancers.

## Acknowledgements

We thank the NYU Genome Technology Center (RRID: SCR_017929) and the NYU Cytometry and Cell Sorting Laboratory (RRID: SCR_019179).

---

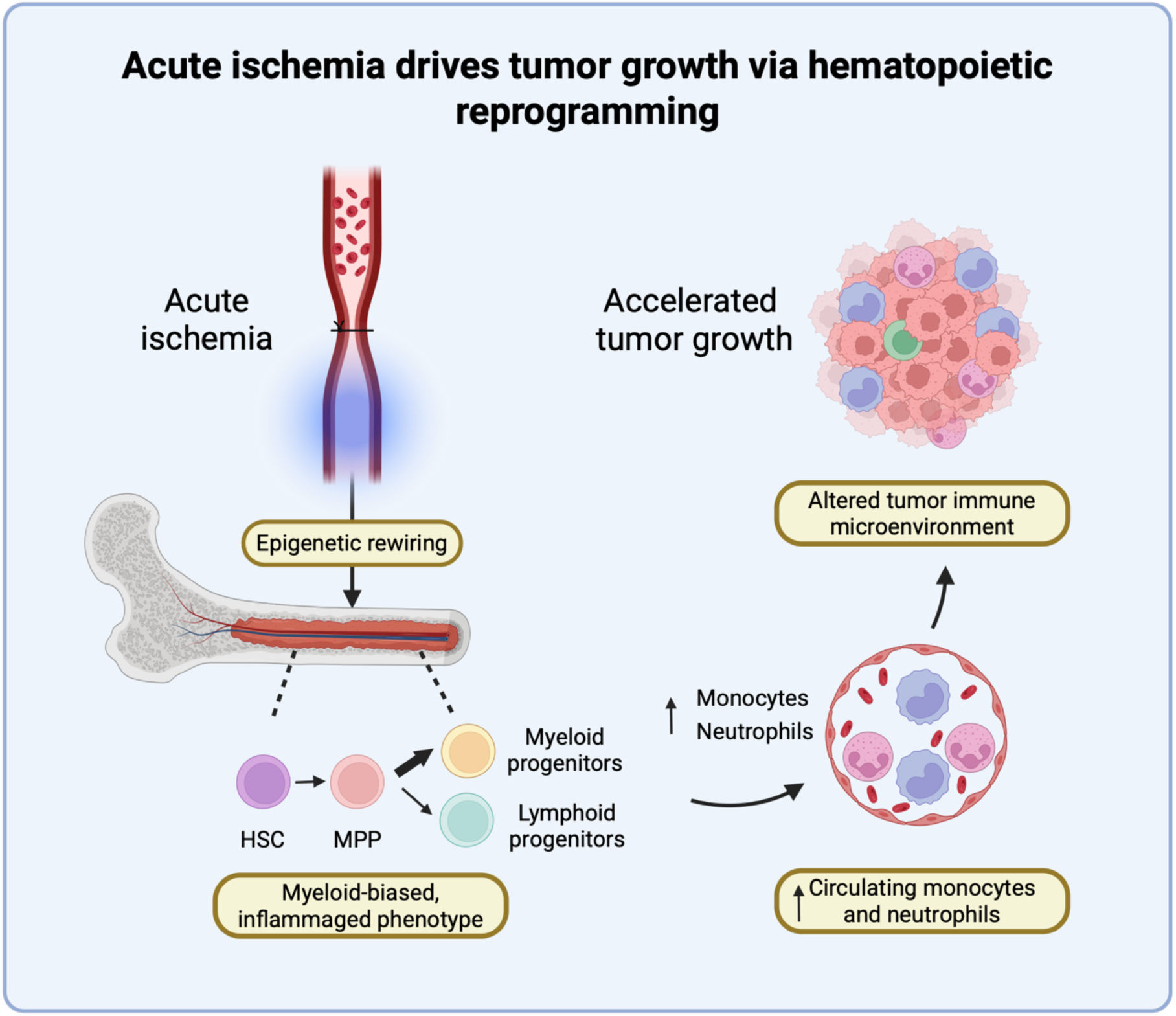
Central Illustration. Peripheral ischemic injury accelerates cancer outgrowth through maladaptive reprogramming of myelopoiesis. Ischemia induces inflammatory and epigenetic alterations in bone marrow stem and progenitor cells that result in myeloid bias and an immunosuppressive tumor microenvironment.

