## Supplemental File for "Ischemic Injury Drives Tumor Growth via Accelerated Hematopoietic Aging"

**Supplemental Figures 1-11**
**Supplemental Figure Legends**
**Supplemental Methods**

#### Figure S1

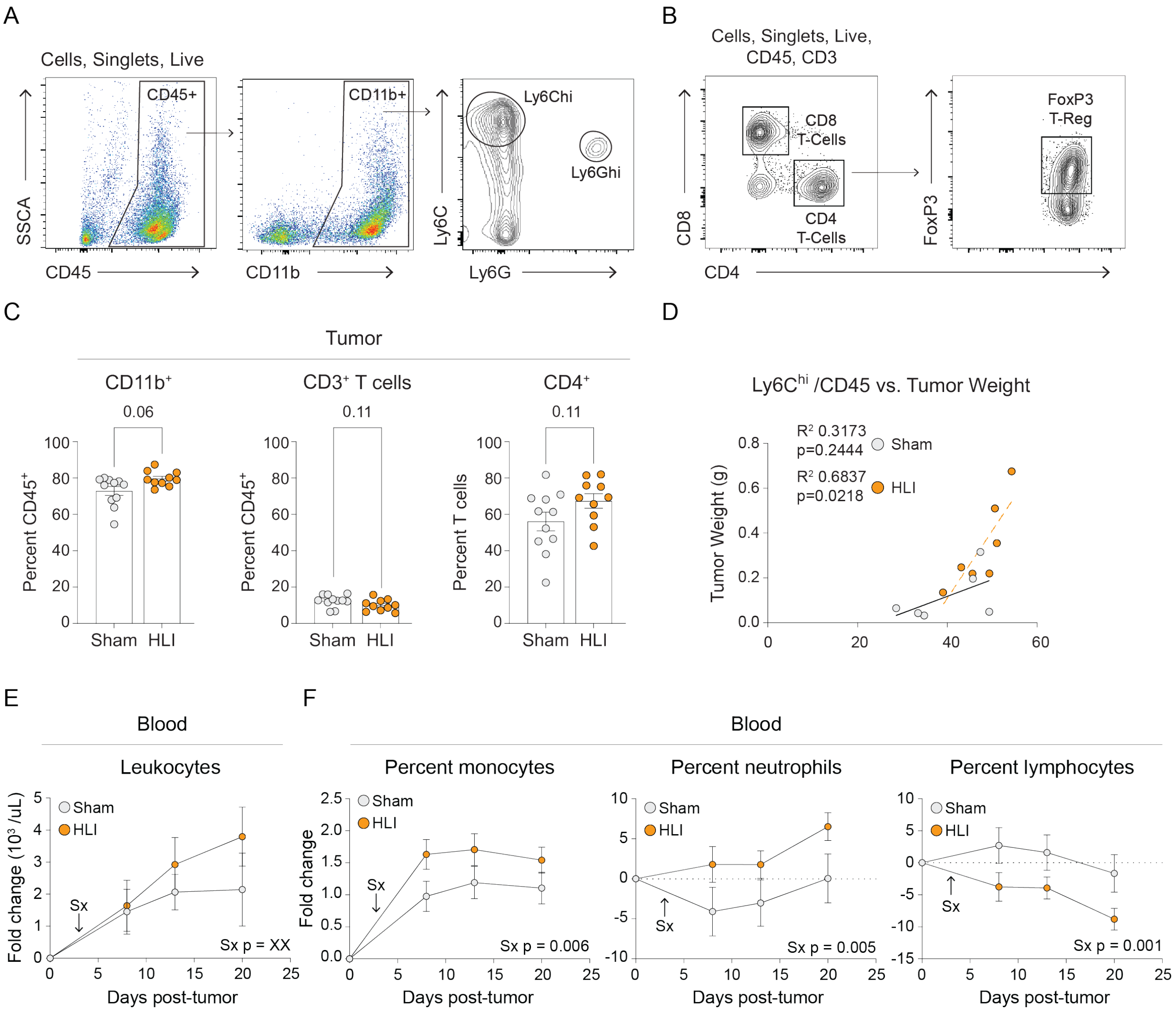


**Supplemental Figure 1: Hindlimb ischemia alters immune cells in the tumor and circulation.** (A,B) Flow cytometry gating strategy for (A) myeloid and (B) T cells. (C) $CD11b^+$ myeloid cells, $CD3^+$ and $CD4^+$ T cells in the tumor 21 days after surgery. (D) Simple linear regression analysis of the proportion of $CD11b^+$ $Ly6C^{hi}/CD45^+$ cells in the tumor vs. tumor weight. (E) Absolute number of circulating leukocytes over time. (F) Relative frequency of leukocytes in the circulation over time. FC: fold change. Student's t-test (C), simple linear regression (D), or Two-way ANOVA (E, F). Mean ± SEM.

### Figure S2

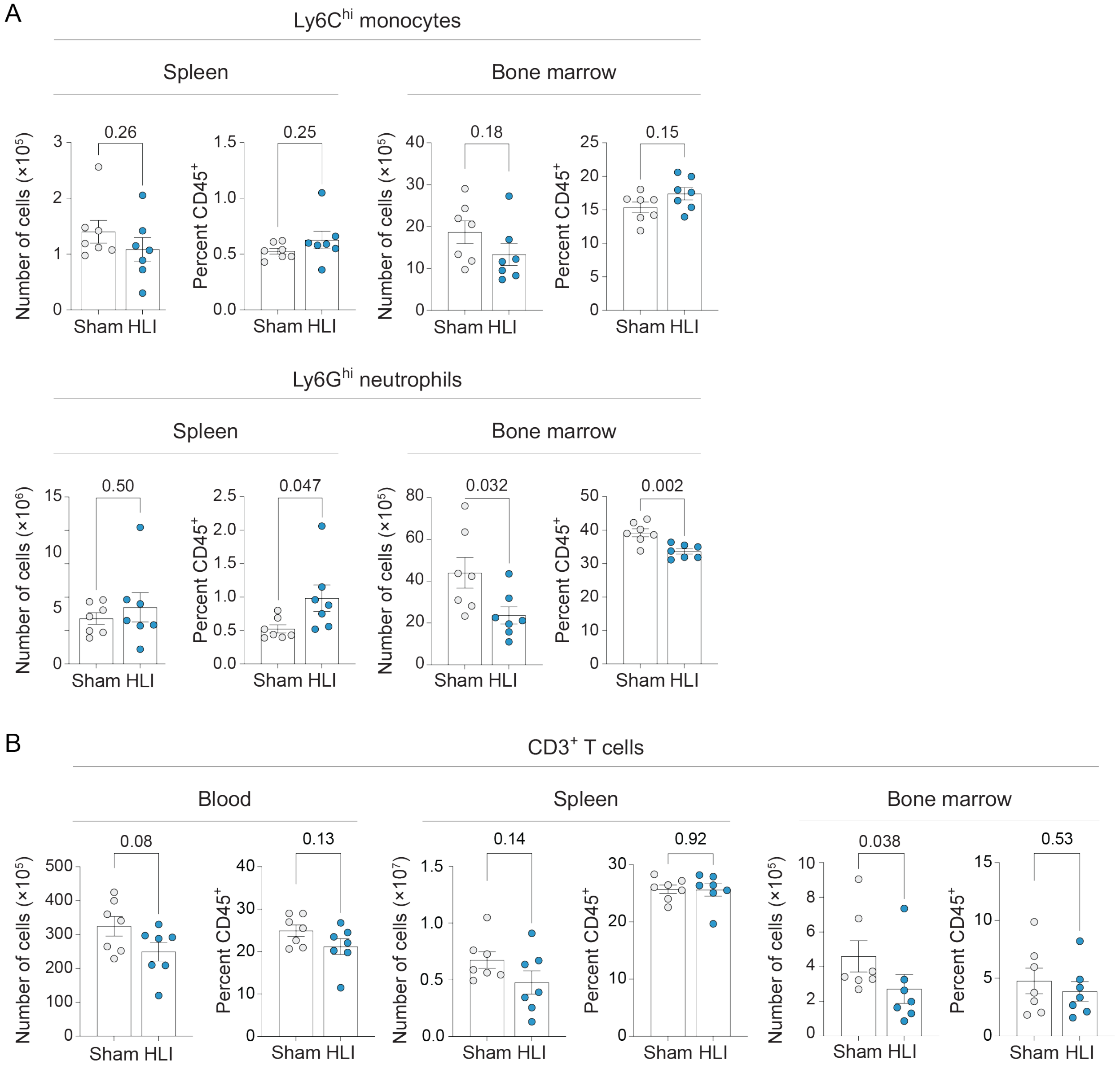


**Supplemental Figure 2: Mature immune cells are altered in the spleen and bone marrow after HLI.** (A,B) Absolute number and relative frequency of (A) of CD11b$^+$ Ly6C$^{hi}$ monocytes and (B) of CD11b$^+$ Ly6G$^{hi}$ neutrophils in the spleen (left) and bone marrow (right) two days after HLI. (C) Absolute number and relative frequency of CD3$^+$ T cells in the blood, spleen, and bone marrow in mice exposed to HLI surgery or sham surgery. Student's t-test (A,B). Mean ± SEM.

#### Figure S3

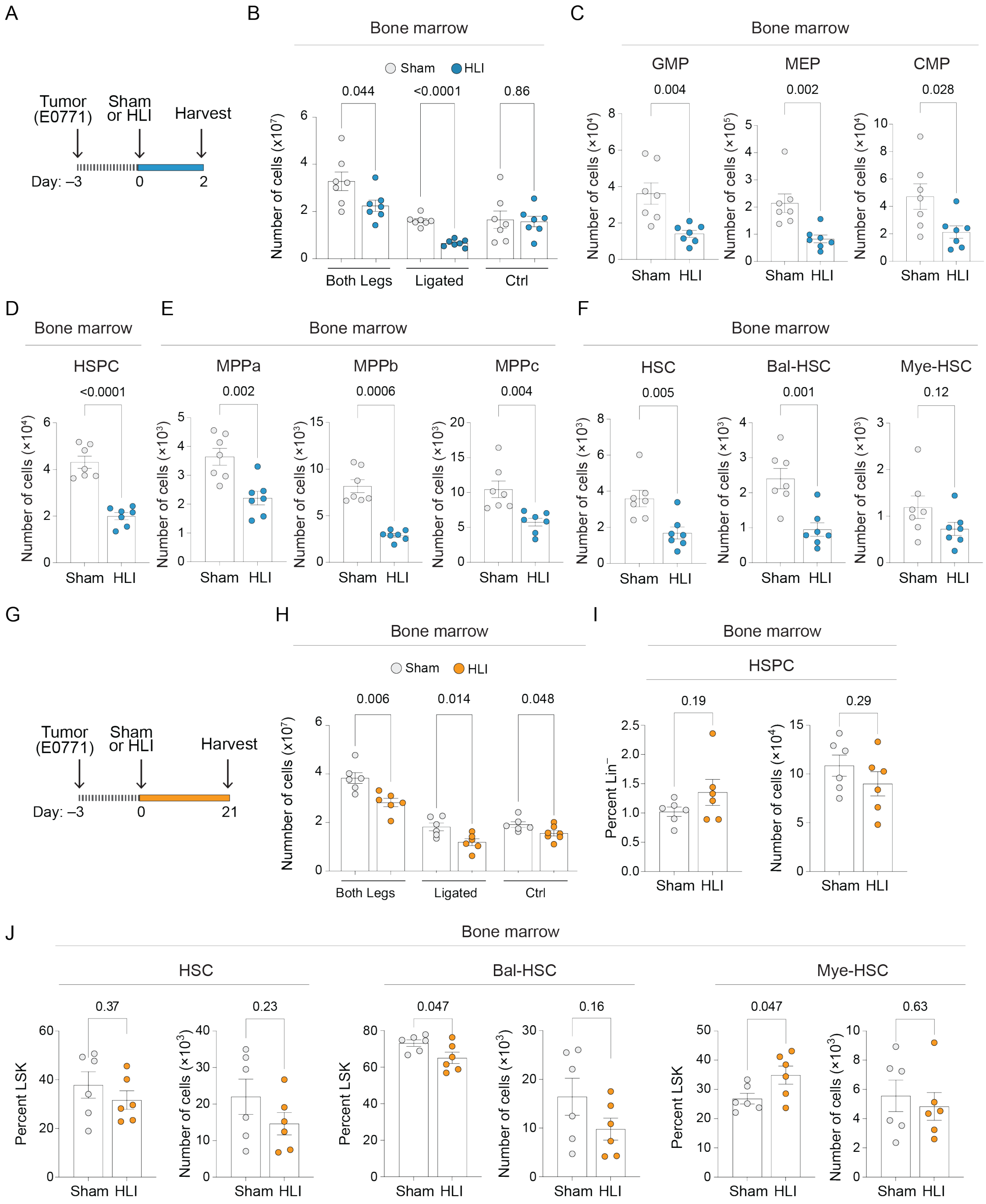


**Supplemental Figure 3: HLI depletes cells in the bone marrow.** (A) Experimental design (2 days post-surgery). (B) Number of cells in the bone marrow. (C-F) Number of progenitors in the ligated femur and tibia including (C) GMP, MEP, CMP, (D) HSPC, (E) MPPa, MPPb, MPPc, and (F) total, balanced and myeloid-biased HSCs. (G) Experimental design (21 days post-surgery). (H) Number of cells in the bone marrow. (I-J) Absolute number and relative frequency of (I) HSPC and (J) total, balanced, and myeloid-biased HSCs. Student's t-test (B, C, D, E, F, H, I, J) or Mann-Whitney U-test (MPPb). Mean ± SEM.

#### Figure S4

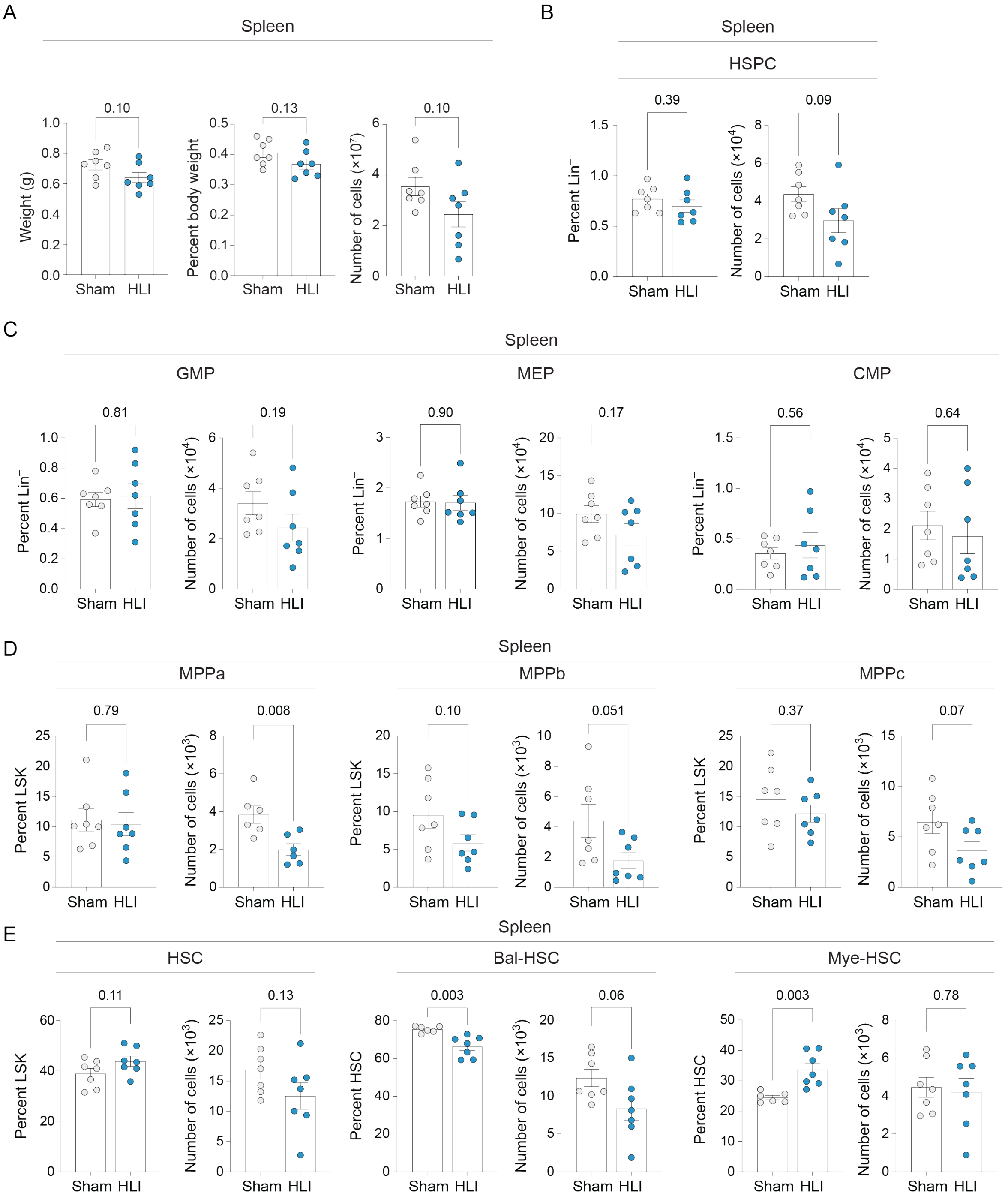


**Supplemental Figure 4: HLI induces a myeloid bias in hematopoietic progenitors in the spleen** (A) Spleen weight and cellularity two days after HLI or sham surgery. (B-E) Absolute number and relative frequency of progenitors in the spleen including (B) HSPC, (C) GMP, MEP, CMP, (D) MPPa, MPPb, MPPc, and (E) total, balanced and myeloid-biased HSCs two days after HLI or sham. Student's t-test (A,B,C,D,E) or Mann-Whitney U-test (MPPb). Mean ± SEM.

#### Figure S5

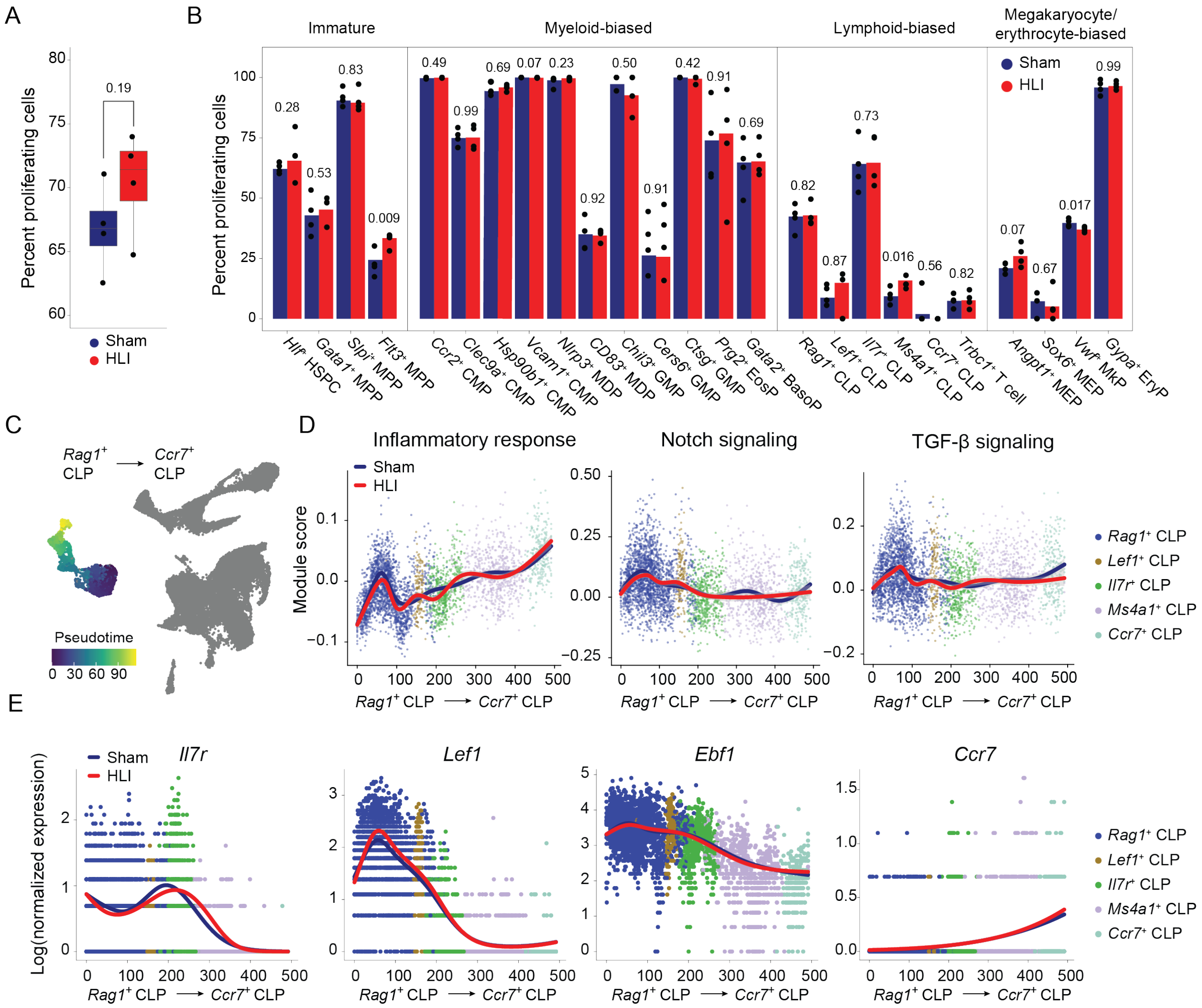


**Supplemental Figure 5: scRNA-sequencing indicates HLI induces lymphoid dysfunction**
(A-B) Relative frequency of proliferating bone marrow progenitor cells in mice exposed to HLI versus sham surgery. (A) Total and (B) separated by island: immature, myeloid-biased, lymphoid-biased, and megakaryocyte/erythrocyte-biased. (C-E) scRNA-seq indicates a single lymphoid progenitor trajectory. (C) UMAP trajectory, (D) hallmark gene set enrichment analysis, and (E) transcript expression of lymphocyte marker genes along the *Ccr7*$^{+}$ trajectory.

**Figure S6**

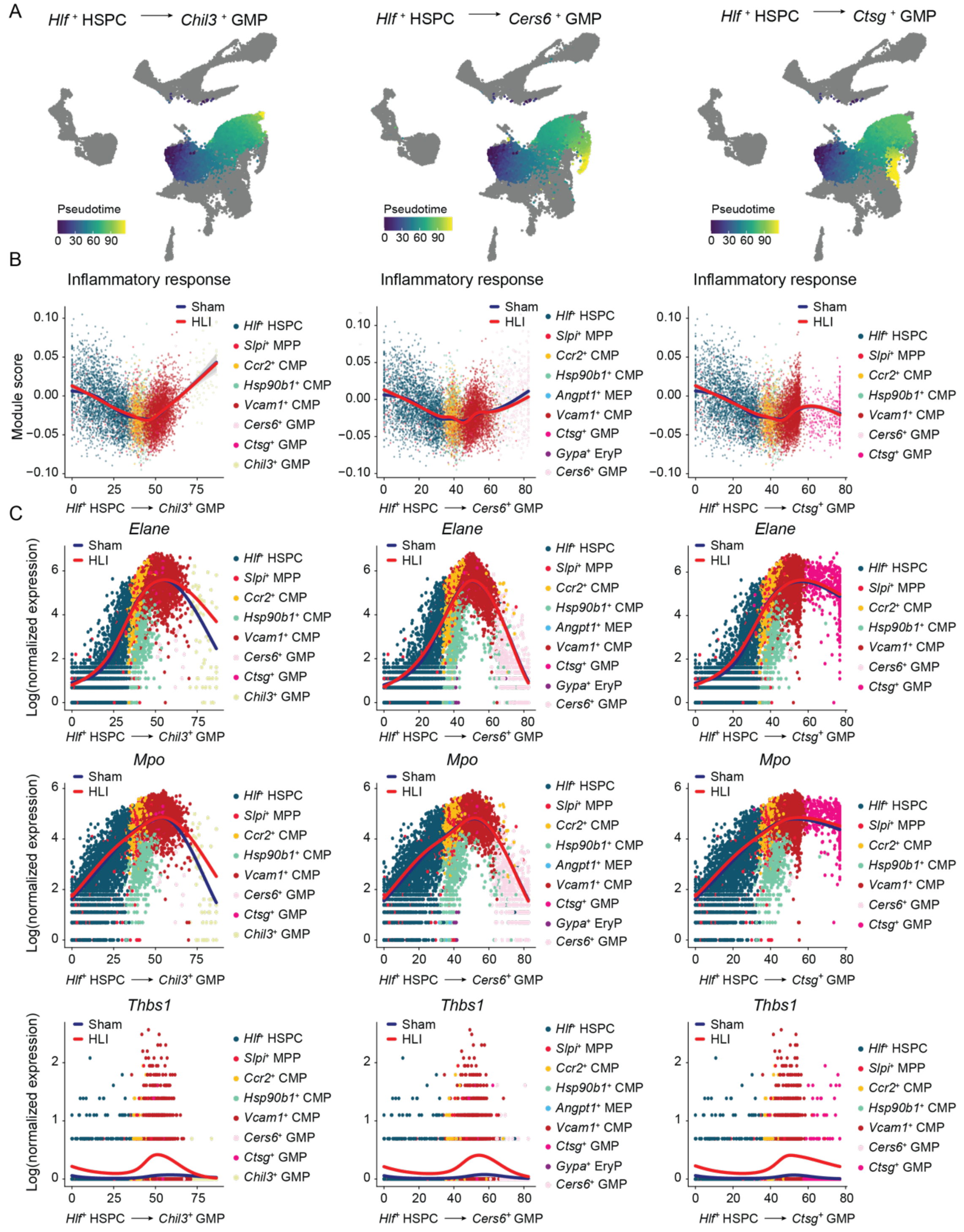


**Supplemental Figure 6: scRNA-sequencing identifies three GMP trajectories in the bone marrow.** (A) UMAP trajectories, (B) hallmark gene set enrichment analysis, and (C) transcript expression of GMP marker genes along the *Chil3*⁺ (left), *Cers6*⁺ (center), and *Ctsg*⁺ (right) trajectories.

### Figure S7

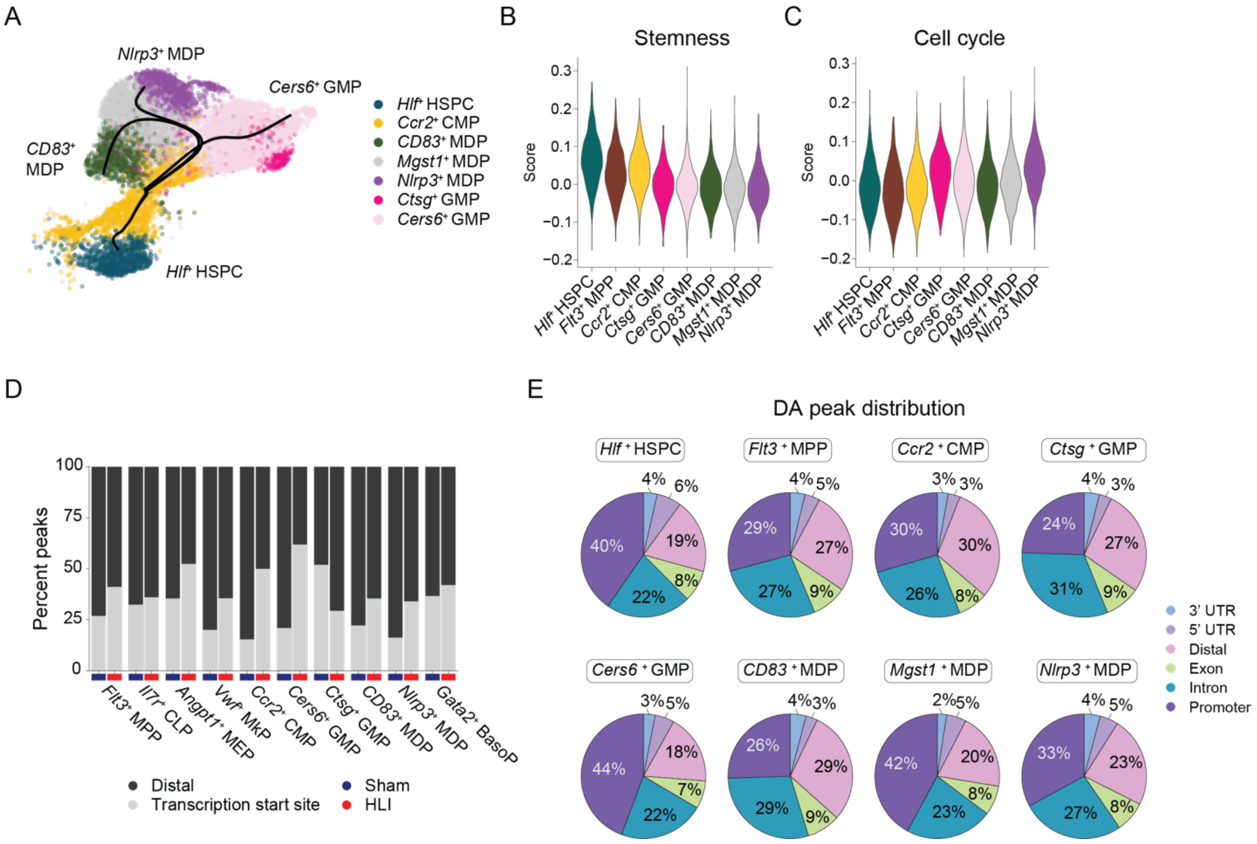


**Supplemental Figure 7: Myeloid differentiation trajectories and differential chromatin accessibility in myeloid progenitors after HLI** (A) UMAP showing myeloid differentiation trajectories. (B) Violin plots demonstrating stemness scores across myeloid progenitor populations. (C) Violin plots demonstrating cell cycle scores across myeloid progenitor populations. (D) Percent peaks distal and within transcription start site in myeloid progenitor populations. (E) Percentage peak distribution by location in each myeloid progenitor population. DA, differentially accessible

### Figure S8

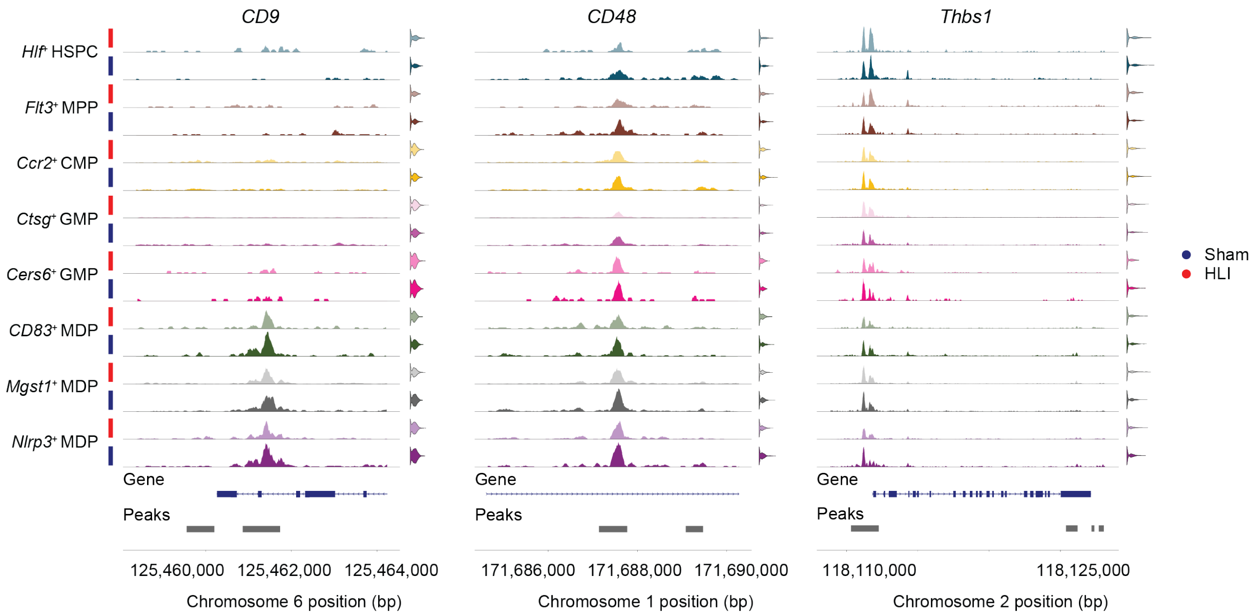


**Supplemental Figure 8: Chromatin accessibility peaks in myeloid progenitors after HLI**
Tracks displaying sequencing read coverage across indicated loci, stratified by treatment.

### Figure S9

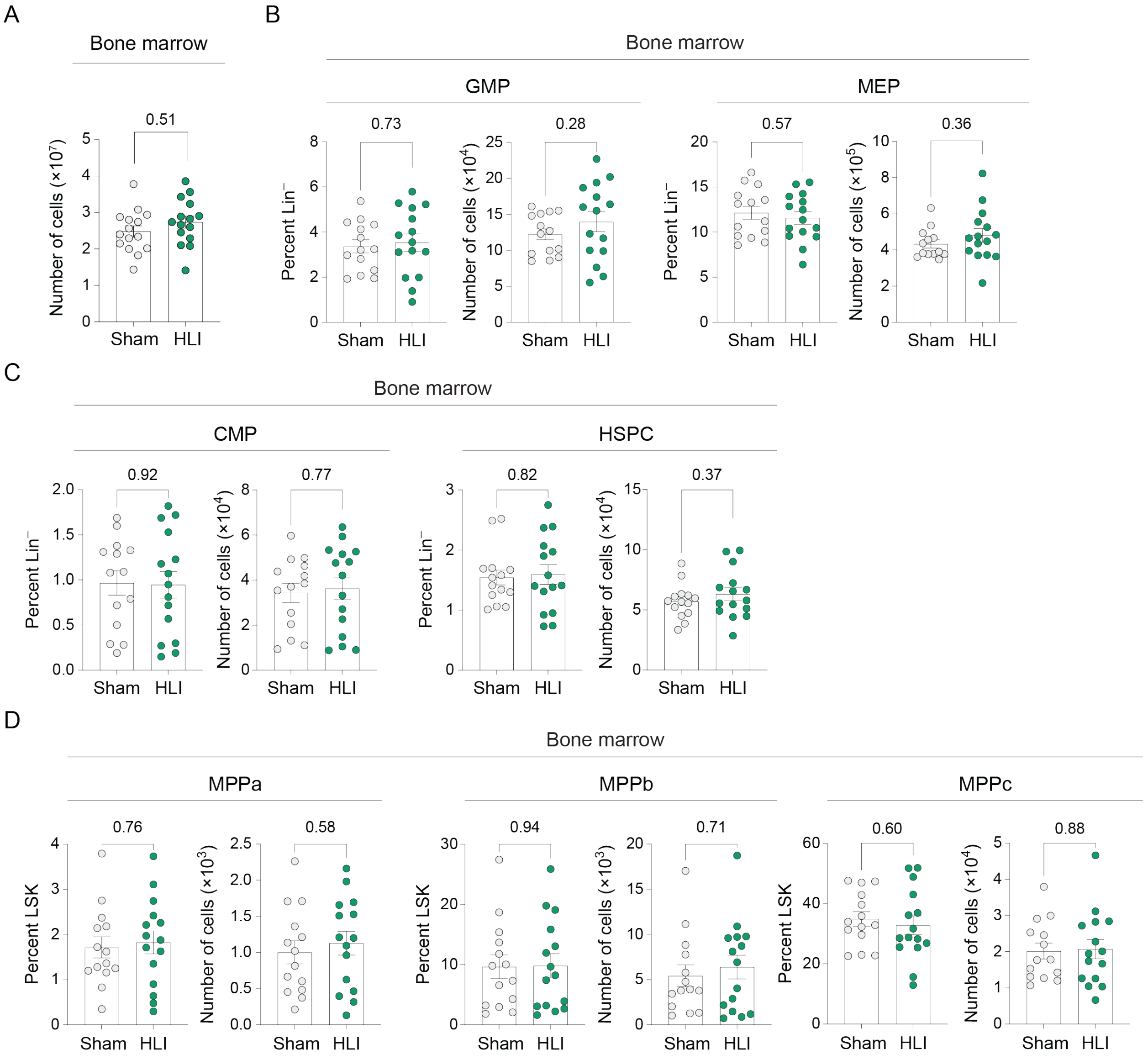


**Supplemental Figure 9: Bone marrow characterization of HLI- and sham-bone marrow recipients bearing E0771 tumors.** (A) Number of cells in the bone marrow of E0771 tumor-bearing mice that received HLI versus sham bone marrow at study endpoint. (B-D) Relative frequency and absolute number of progenitors in the bone marrow of mice that received HLI versus sham bone marrow: (B) GMP, MEP, (C) CMP, HSPC, and (D) MPPa, MPPb, MPPc. Student's t-test (A,B,C,D) or Mann-Whitney U-test (MPPb). Mean ± SEM.

#### Figure S10

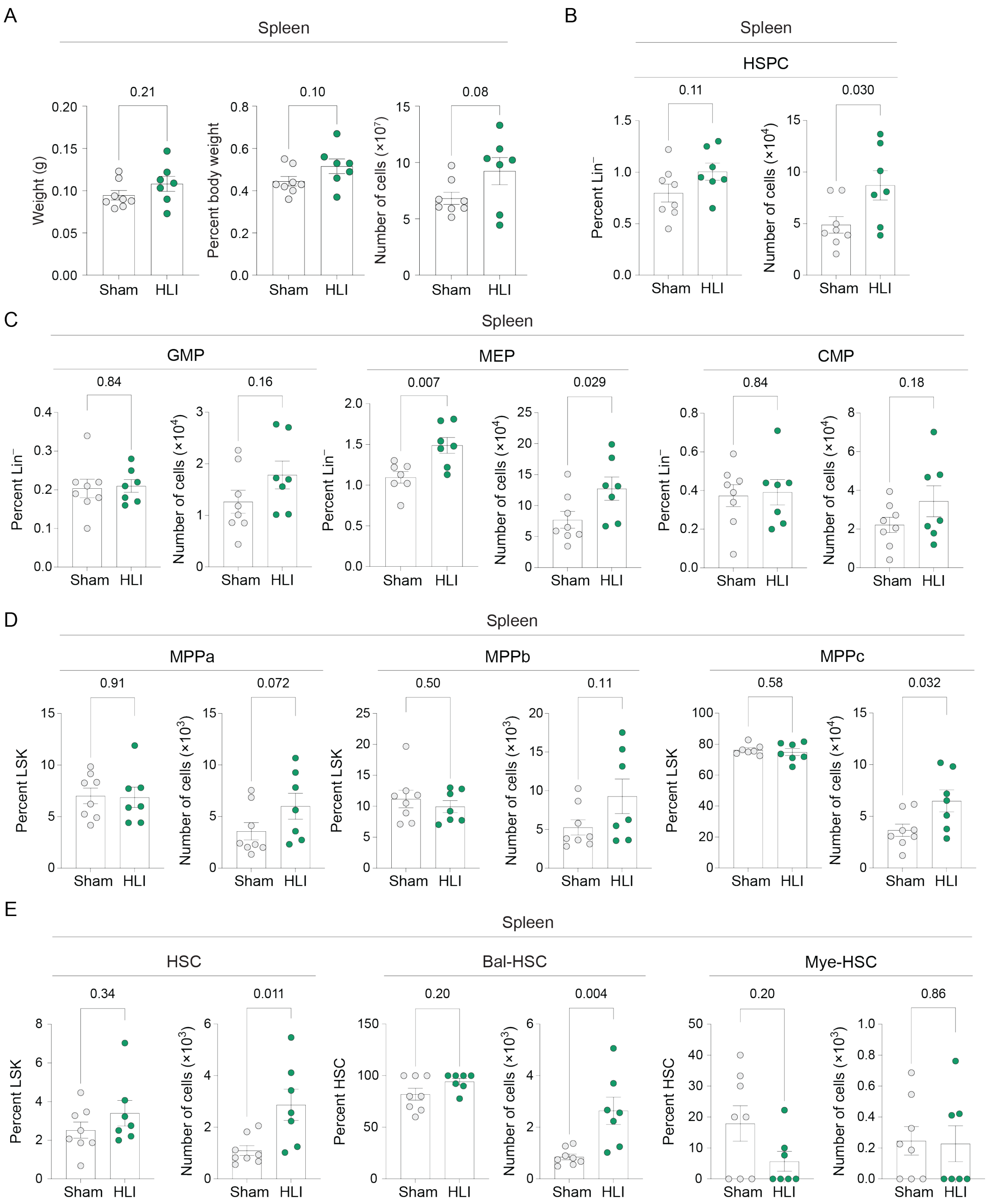


**Supplemental Figure 10: Characterization of spleen hematopoietic progenitors in HLI- and sham-bone marrow recipients bearing E0771 tumors.** (A) Spleen weight and cellularity at study endpoint. (B-D) Absolute number and relative frequency of splenic progenitors including (C) HSPC, GMP, MEP, CMP, (C) MPPa, MPPb, MPPc, and (D) HSC and balanced and myeloid-biased HSCs in HLI or sham bone marrow recipient mice. Student's t-test (A,B,C,D) or Mann-Whitney U-test (MPPb). Mean ± SEM.

### Figure S11

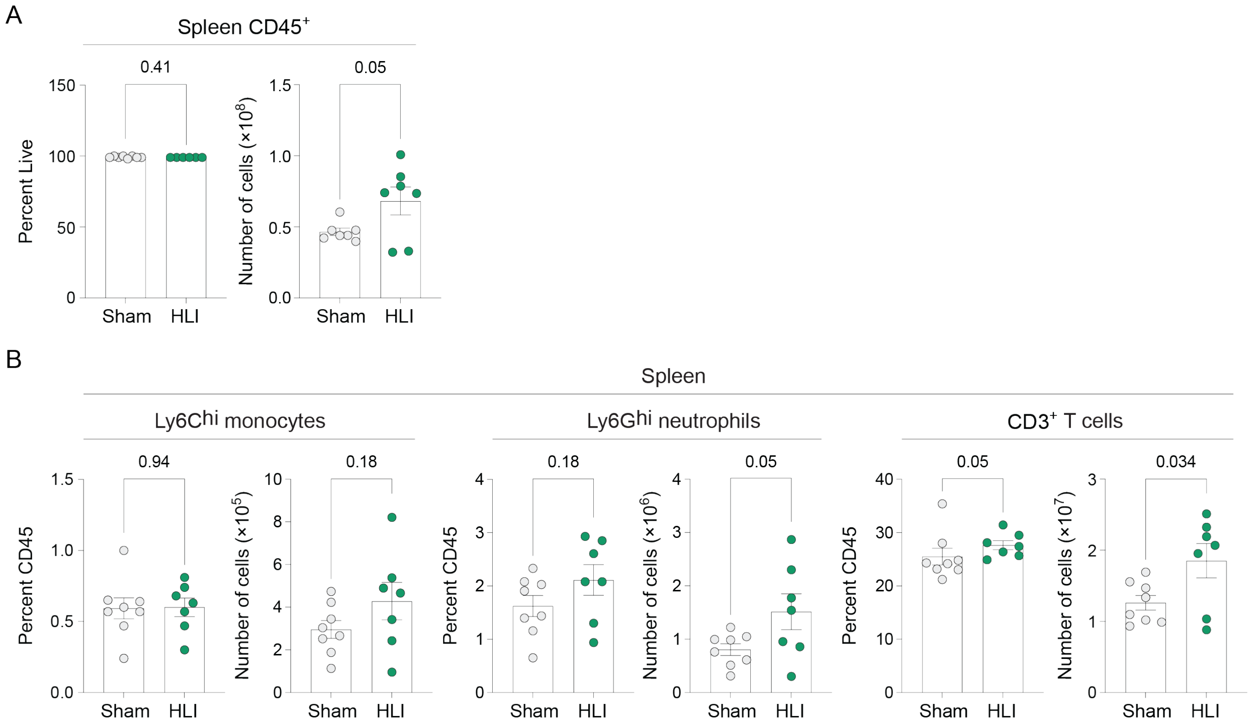


**Supplemental Figure 11: Immune cell profile of the spleen of bone marrow recipient mice**
(A) Relative frequency and absolute number of $CD45^+$ leukocytes in the spleens of E0771 tumor-bearing mice that received HLI versus sham bone marrow at study endpoint. (B) Relative frequency of $CD11b^+$ $Ly6C^{hi}$ monocytes, $CD11b^+$ $Ly6G^{hi}$ neutrophils, and $CD3^+$ T cells in the spleen. Student's t-test (A,B) or Mann-Whitney U-test (CD3). Mean ± SEM.

**Supplemental Methods:**

*Laser doppler acquisition and perfusion analysis*

Perfusion in the hind limb was measured using high-resolution laser doppler imaging (moorLD12-HIR, moor Instruments). Mice were anesthetized, maintained at a flow rate of 1.5% isoflurane and $O_2$, and hindlimbs were shaved to expose the dermis. Mice were kept on a heating pad to maintain their body temperature. Measurements were recorded directly after (day 0) as well as on days 8 and 17 following femoral artery ligation or sham surgery. Scans were analyzed using moorLD12-HIR software. Blood flow was assessed as mean flux by drawing ROIs of identical size in the ischemic and non-ischemic legs. Perfusion was quantified as the ratio of mean flux in the left limb (ligated or sham-ligated) to the right limb (contralateral control).

*Flow cytometry*

Flow cytometry analysis was performed on tumors, bone marrow and blood as described in supplemental methods. For flow cytometry, tumors were weighed, rinsed in MACS buffer, digested with MACS Miltenyi tumor kit (Miltenyi Biotec, 130-096-730) and dissociated with the gentleMACS Dissociator (Miltenyi, 130-093-235) according to manufacturer's instructions. Bone marrow was collected from the long bones in the left and right legs, kept separate, and red blood cells (RBS) were lysed with ACK lysis buffer (Fisher, A1049201). Spleens were weighed, rinsed in MACS buffer, and passed through a 70μm filter. RBC were lysed with ACK lysis buffer (Fisher, A1049201), passed through a 40um filter. Cells were counted using ThermoFisher Cell Countess 2.

Blood (~30μL) was processed with ACK lysis buffer (Fisher, A1049201) and fixable live/dead staining performed in PBS for 30 minutes (eFluor 780 - EBioscience cat. 65-0865-14, eFluor 506 - eBioscience cat. 65-0866-14). All other extracellular stains were performed in MACS/FACS buffer. Cells were incubated with Fc block (cat) for 5 minutes, except bone marrow progenitors. Intracellular stains were performed with True Nuclear Transcription Factor Buffer Set (BioLegend 424401) according to manufacturer's protocol. Cells were run on a MACSQuant 10 or 16 machine and data analyzed using FlowJo software. Single, live, cells were gated by SSC and FSC to exclude debris and doublets and by negative staining of live/dead marker.

Mobilization of cells from the spleen and bone marrow was assessed as the ratio of $CD11b^{+}$ $Ly6C^{hi}$ $Ly6G^{lo}$ monocytes or $CD11b^{+}$ $Ly6G^{hi}$ $Ly6C^{int}$ neutrophils in the blood and the tissue. Frequency analyses were performed by dividing the cell count for each population over the total bone marrow count.

Reagent Table 1:

| **Antibody – Fluorophore** | **Company** | **Clone** | **Catalog** |
|---|---|---|---|
| CD34 – eF450 | Inv | RAM34 | 48-0341-82 |
| CD34 – AF647 | BD | RAM34 | 560230 |
| CD62P – SB600 | Inv | Psel.KO2.3 | 63-0626-82 |
| CD117 – BV650 | Inv | 2B8 | 416-1171-82 |
| CD117 – APC-Cy7 | BL | 2B8 | 105826 |
| CD135 (Flt3) – PE | BL | A2F10 | 135306 |
| Sca1 – PacBlue | BL | D7 | 108120 |
| Sca1 – PE-TxRed | BL | D7 | 108138 |
| CD150 – BV605 | BL | TC15-12F12.2 | 115927 |

| | | | |
|---|---|---|---|
| CD150 – PerCP-eF710 | Inv | mShad150 | 46-1502-82 |
| CD16/32 – PE-Cy7 | BL | 93 | 101318 |
| CD48 – PE-Cy5 | BL | HM48-1 | 103420 |
| CD48 – AF647 | Inv | HM48-1 | 17-0481-82 |
| Lin – AF700 | BL | Cocktail | 133313 |
| Lin – FITC | BL | Cocktail | 133302 |
| CD45.2 | BL | 104 | 109831 |
| CD45.1 | BL | A20 | 110718 |
| Ly6C – BV421 | BL | HK1.4 | 128031 |
| Ly6C – APC-Cy7 | BL | HK1.4 | 128026 |
| CD11b – PE-Cy7 | BL | M1/70 | 101216 |
| F4/80 – FITC | BL | BM8 | 123108 |
| CD45 – FITC | BL | | 103108 |
| CD45 – PerCP-Cy5.5 | BL | 30-F11 | 103132 |
| CD45 – BV510 | BL | 30-F11 | 103138 |
| IA/IE – AF700 | BL | M5/114.15.2 | 107622 |
| Ly6G – PE | BL | 1A8 | 127608 |
| FoxP3 – BV421 | BL | MF-14 | 126419 |
| CD8 – FITC | BL | 53-6.7 | 100706 |
| CD3 – PerCP-Cy5.5 | BL | 17A2 | 100218 |
| CD3 – FITC | BL | 17A2 | 100204 |
| CD3 – AF700 | BL | 17A2 | 100216 |
| CD4 – PE-Cy7 | BL | GK1.5 | 100422 |

*Single Nucleus Multiome ATAC-sequencing and Gene Expression*

For snATACseq with gene expression ("snMultiomics"), nuclei were extracted from the sorted cells using a modified nuclei isolation protocol. Cells were spun down and washed two times with wash buffer (PBS + 0.04% BSA) at 4˚C. Supernatant was aspirated and cells were lysed by pipetting up and down 10 times in 100µL of chilled lysis buffer (10mM Tris-HCl pH 7.4, 10mM NaCl, 3mM MgCl2, 0.1% IGEPAL CA-630, 0.1% Tween-20, 0.01% Digitonin, 1% BSA, 1mM DTT 1U/µL RNase inhibitor). Immediately after, 1000µL of nuclei extraction wash buffer (10mM Tris-HCl pH 7.4, 10mM NaCl, 3mM MgCl2, 0.1% Tween-20, 1% BSA) was added to the sample and the nuclei were pelleted. This was repeated two more times for a total of three washes. The nuclei were then pooled and snRNA-seq and snATAC-seq libraries were constructed using Chromium Next GEM Single Cell Multiome ATAC + Gene Expression Reagent Kit (10x Genomics, PN-10000285) according to the manufacturer's protocol. Briefly, 10K nuclei in 1x nuclei buffer were loaded with a target output of ~6,000. Single nuclei were encapsulated into emulsion droplets using Chromium Controller (10x Genomics). Transposition, reverse transcription, and library preparation were performed on C1000 Touch Thermal cycler with 96-Deep Well Reaction Module (Bio-Rad). Amplified cDNA was evaluated on an Agilent BioAnalyzer 2100 using a High Sensitivity DNA Kit (Agilent Technologies, PN5067-4626) and final libraries on an Agilent TapeStation 4200 using High Sensitivity D1000 ScreenTape (Agilent Technologies, PN5067-5584). Individual libraries were normalized to 2nM and pooled for sequencing. Pools were sequenced with 10B 200 cycle run kits (50bp Read1, 10bp Index1 24bp Index2 and 90bp Read2) on the NovaSeq X+ Sequencing System (Illumina) and analyzed using CellRanger software.

Reagent Table 2:

| Item | Manufacturer | Catalog Number |
|---|---|---|
| Tween-20 | BioRad | 1662404 |
| RNase inhibitor | Sigma | 3335399001 |
| AOPI | Revvity | CS2-0106 |
| RNase Free PBS | Invitrogen | AM9624 |
| NaCl (5 M), RNase-free | Fisher | AM9760G |
| IGEPAL CA-630 | Sigma | I8896-50ML |
| Digitonin | Sigma | D141-100MG |
| Magnesium Chloride Solution, 1M | Sigma | M1028 |
| Tris-HCl | ThermoFisher | 15567027 |
| 20X Nuclei Buffer | 10x Genomics | 2000207 |

*scRNAseq*

Sorted cells were processed for scRNAseq and hashtagged using Type A oligo-tagged antibodies (Biolegend TotalSeq™-A0301/A0304 anti-mouse Hashtags, 155801-09). After staining, cell number and viability were assessed using a TC20 Cell Automated Cell Counter (BioRad). To reduce experimental batch effects, specimens from the two experimental groups were pooled into two preps (2 sham + 2 HLI per prep) after sorting and hashtagging. Briefly, single cell suspensions were partitioned into PIPs (PIPseq Epitope Sequencing Kit, T20 V4.0PLUS) with subsequent cDNA generation and final library preparation following manufacturer's instructions. The two preps contained 54,000 (sample A) and 42,000 (sample B) viable cells. cDNA quality and concentration were evaluated on a BioAnalyzer 2100 using a High Sensitivity DNA Kit (Agilent Technologies). Final gene expression and HTO libraries were amplified 10 and 8 cycles respectively and visualized on an Agilent TapeStation 4200 using High

Sensitivity D1000 ScreenTape (Agilent Technologies). Sequencing was performed using a NovaSeq X+ 10B 200 Cycle Flowcell (Illumina).

*Data analysis: Multiome*

ATAC-seq reads and gene expression from the 2 samples were initially processed through the 10x Genomics software Cell Ranger ARC (v2.0.0) for barcode demultiplexing, cell calling, genome alignment (GRCm38), RNA-seq feature counting (GENCODE vM23), and an initial round of ATAC-seq peak calling. RNA-seq and ATAC-seq count data were further processed using the R packages Signac (v1.13.0)[66] and Seurat (v4.4.0).[67,68] Low quality barcodes/cells were removed if they contained less than 680 RNA UMIs, percent mitochondrial genome mapping UMIs greater than 73%, ATAC UMIs less than 2000, ATAC nucleosome signal greater than 1.2, ATAC TSS enrichment less than 4, percent ATAC fragments in peaks less than 40%, and percent ATAC fragments in TSS regions less than 27%. Mitochondrial genes were then removed from the RNA count matrix.

Following filtering, peaks were called again for each sample using MACS2 within the Signac function CallPeaks(). The data from both samples were then merged. For the ATACseq data this necessitated merging the peak sets from both samples into a unified peak set, which was accomplished via Signac's reduce() function. Merged peaks that were less than 20bp or greater than 10,000 bp in width were removed. Fragments were then quantified within each peak and cell via Signac's FeatureMatrix() function. At this point, the merged RNA-seq and ATAC-seq data were processed separately.

For the ATAC-seq data, Signac's FindTopFeatures() was run to select the most variable peaks. To reduce dimensionality we employed latent semantic indexing (LSI) via Signac's

RunTFIDF() function followed by RunSVD(). A shared nearest neighbor (SNN) graph was constructed from the LSI dimensions 2-30 using Seurat's FindNeighbors() function. To cluster cells, the leiden algorithm was then applied via Seurat's FindClusters() function. Of the 15 clusters identified, one showed distinctly poor ATAC and RNA quality metrics, e.g. very few genes detected per cell and a low number of ATAC UMIs in peaks. We determined these cells were low quality and removed them from the dataset.

For the RNA-seq data, normalization was performed using Seurat's SCTransform() function. Dimensionality reduction was performed using principal component analysis (PCA) via Seurat's RunPCA() function. The first 30 PCA dimensions were used to construct a SNN graph upon which the Leiden algorithm was applied as mentioned above. It became evident that a substantial batch effect was present in the RNA data driven in part by a large difference in mitochondrial gene counts. To mitigate this, we employed Seurat's canonical correlation analysis (CCA) based integration method. To this end the Seurat functions, SelectIntegrationFeatures(), PrepSCTIntegration(), FindIntegrationAnchors, and IntegrateData() were run in the order mentioned. Dimensionality reduction and clustering were performed on the integrated data as mentioned above for the unintegrated RNA data.

To define cell states using both modalities, Seurat's weighted nearest neighbor analysis (WNN) was applied. To this end, Seurat's FindMultiModalNeighbors() function was run using dimensions 1-30 of the integrated RNA PCA and dimensions 2-30 of the ATAC LSI. The leiden algorithm was then applied to the resulting weighted nearest neighbor graph via Seurat's FindClusters() function. UMAP was run on the WNN dimensions 1-30 via Seurat's RunUMAP() function.

To identify accessible regions that may be specific to smaller populations of cells, peak calling was run again for each of the 22 WNN clusters using MACS2 as described above with the group.by parameter set to the WNN clusters. Fragments were then quantified within each peak and cell via Signac's FeatureMatrix() function. To use this more nuanced peak set in the identification of different cell states, the WNN and clustering procedure described above was repeated using 22 clusters.

To aid in cell state annotation, gene activity scores were calculated using Signac's GeneActivity() function. Cluster defining RNA and gene activity signatures were calculated using wilcoxauc() from the R package presto. Cluster defining ATAC peak signatures were identified using Seurat's FindAllMarkers() function with parameters with parameters: only.pos = T, logfc.threshold = 1.5, test.use = 'LR', and latent.vars = 'nCount_peaks'. Cluster identification and labaling was performed by canonical marker gene expression computed via the pseudobulk method developed by Squair et al.[69] Multiplets, low-quality cells, and terminally differentiated cells were filtered out from the dataset. Proliferation and stem scores were computed on gene activities using the AddModuleScore() function from Seurat. The stem gene list was obtained from Giladi et al.[30] The cell cycle gene list used was a concatenation of Seurat's S and G2M cc.genes list.

Trajectory analysis was performed using the R package Slingshot.[31] ATAC LSI dimensions 2-10 were used with the starting cluster set to HSPC. CLP clusters were processed using the omega parameter set to TRUE to allow for disconnected trajectories. Principal curves were embedded in the 2 WNN UMAP dimensions using Slingshots embedCurves(), followed by slingCurves().

Differential accessibility analysis was performed between sham and HLI cells within each cluster using Seurat's FindAllMarkers() function with parameters logfc.threshold = 0.5, test.use = 'LR', latent.vars = 'nCount_peaks', min.cells.group = 3, min.cells.feature = 3, min.pct = 0.1. To link peaks to genes, the R package rGREAT was used with default parameters on the whole peakset. Peaks were classified as transcription start site (TSS) if they were < 3kb up or downstream from an annotated TSS, otherwise, they were categorized as distal. rGREAT was also run on subsets of peaks for functional enrichment. For more granular peak location assignments (Fig S7D), clusterprofiler was used with default parameters. Transcription factor activities were estimated using chromVAR[43] via Signac's RunChromVAR() function. To compare transcription activities between Sham and HLI cells within each cluster, Seurat's FindMarkers() function was used.

*Data analysis: scRNA-seq*

Two datasets (A&B) each containing 4 hashtagged samples were run through the Pipseeker software from Fluent Biosciences (v.3.0.5). Pipseeker performs alignment (GRCm39, GENCODE vM29), filtering, barcode counting and unique molecular identifier (UMI) counting of RNA and hash-tagged oligo (HTO) sequencing reads from fastq files. The Seurat package (v.4.4.0) in R (v.4.1.2) was used for sample processing and analysis. Pipseeker count matrices corresponding to a cell calling sensitivity of 5 were analyzed. Only genes with 1 or more counts in at least 0.1% of barcodes were retained. HTO demultiplexing was performed with Seurat by first normalizing the HTO counts using NormalizeData() with normalization.method="CLR". Singlet, doublet, and negative barcodes were marked by running HTODemux() with positive.quantile=0.99. Thresholds were manually inspected to insure accurate distinction between negative and positive barcodes for each HTO. RNA data were normalized and scaled using NormalizeData() and ScaleData(), respectively. FindVariableFeautres() was used to identify the 2000 most variable genes. Principal component analysis (PCA) was performed using RunPCA() with npcs = 30. Intra-sample multiplets were detected using recoverDoublets() from the scDblFinder (v.1.8) R package with use.dimred="PCA" and the doublets parameter set to doublets marked by HTODemux() above. Multiplets marked by HTODemux() and recoverDoublets() were removed. Low quality barcodes were then filtered using a combination of genes detected, total UMIs and fraction of UMIs aligning to mitochondrial genes per cell. For sample A, barcodes with percent mitochondrial mapping reads > 5.322%, UMIs per cell > 30000, and genes per cell < $10^{2.8}$ were removed. For sample B, barcodes with percent mitochondrial mapping reads > 5.364%, UMIs per cell > 30000, and genes per cell < 1000 were removed. Samples A and B were then merged yielding 23558 cells. The impact of cell cycle state on clustering was mitigated using a previously published method.[70] Briefly, genes with a Pearson correlation coefficient of at least 0.15 with any one of Ube2c, Hmgb2, Hmgn2, Tuba1b, Ccnb1, Tubb5, Top2a, and Tubb4b we're removed from a duplicated RNA count matrix. This matrix was normalized using SCTransform() with vst.flavor = "v2". PCA was performed on the SCTransform normalized data using RunPCA(). A shared nearest neighbor graph was constructed using the first 30 principal components with FindNeighbors(). FindClusters() was used to cluster the data the Leiden algorithm and a resolution of 0.75. UMAP was computed with the first 30 principal components via RunUMAP(). To identify marker genes of each cluster, a custom pseudobulking script was used to compare cells from each cluster to all other cells via DESeq2. Clusters were annotated using the generated marker gene lists along with canonical cell type markers. To binarize cells based on proliferation state we used a previously published method based on the expression of cell cycle genes defined above.[71] This was used to calculate the fraction of proliferating cells per cluster, condition, and replicate. Trajectory inference and pseudotime ordering was performed using the slingshot() from the Slingshot R package on the first 30 principal components with start.clus = “HSPC”, end.clus = c(“MDP 1”, “GMP 2”, “GMP 3”, “MEP 2”), extend = 'n', stretch = 0, and shrink = 1. Slingshot curves were embedded into the 2 UMAP dimensions using embedCurves() followed by slingCurves(). To identify genes that differ in expression pattern across pseudotime between conditions the R package tradeSeq was used for each lineage of interest.[37] fitGAM() was run using the 5000 most variable genes, nknots = 6, and with the conditions parameter specifying sham and surgery for each cell. Genes with a Benjamini-Hochberg adjusted pval < 0.1 & waldstat > 25 were considered significant. Hallmark pathway scores for each cell were somputed using AddModuleScore() from Seurat.
